# elDORS: An elevated Database Of RNA Sequences

**DOI:** 10.64898/2026.07.10.737016

**Authors:** Nivedita Dutta, Quentin Vicens

## Abstract

Massive and integrated sequence databases have revolutionized computational protein structure prediction. RNA lags due to a lack of consolidated sequence resources. To bridge this gap, we developed elDORS_raw, which comprises up-to-date sequence information, including recent metagenomes and transcriptome sequence data. We optimized an 80% sequence-identity clustered version named elDORS for use with the RNAcmap3 split-strategy for multiple sequence alignment (MSA) and the widely used rMSA pipeline. Our benchmarking demonstrates that elDORS-augmented MSA pipelines match or exceed the alignment depth obtained with massive legacy databases across different queries, including blind CASP challenges, effectively eliminating sequence retrieval failures challenging orphan RNAs. To aid homology searches, predicting RNA properties, training new models, and other downstream tasks, elDORS is freely accessible.

**Graphical Abstract:** 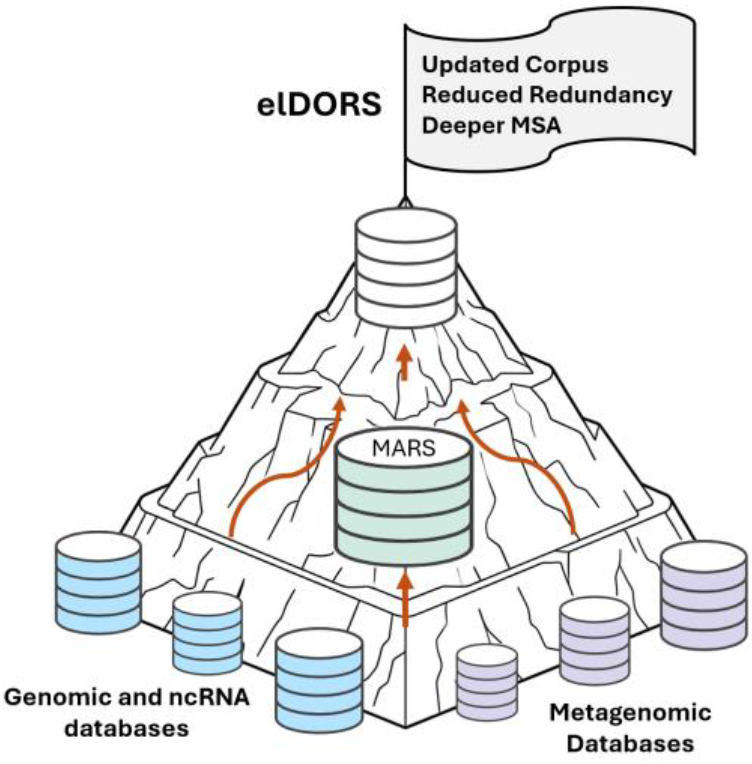

## Introduction

Non-coding and coding RNAs play central roles in catalysis, regulation, and information transfer, among other functions. The associated rapidly growing sequence space captured by high-throughput sequencing is only incompletely represented in current nucleotide repositories. While extensive, existing resources such as NCBI’s nucleotide (nt) collection^1^ and RNAcentral^2,3^ remain limited in depth and diversity for many RNAs, especially metagenomic and non-model transcripts. This limitation is particularly detrimental for high-complexity regions and highly divergent homologs, where sparse sequence data fail to provide the evolutionary intermediates necessary to construct accurate multiple sequence alignments (MSAs). The Master database of all possible RNA Sequences (MARS) ^4^ was originally introduced to address this gap for RNA.

MARS assembled a heterogeneous and large corpus of RNA sequences which, together with the split-search strategy in RNAcmap3 ^4^, enabled far more sensitive homology searches and construction of deeper multiple sequence alignments than previous pipelines, for both Rfam (The RNA families Database) ^5^ and non-Rfam RNAs. These enriched alignments have already translated into improvements in coevolutionary base-pair inference, with RNAcmap3 ^4^ significantly increasing effective sequence depth and improving RNA secondary structure prediction from direct-coupling-analysis (DCA), relative to earlier methods. MSAs generated using RNAcmap3 coupled with the MARS database were utilized for training RNA-MSM, an MSA-based RNA language model (LM) ^6^. RNA-MSM was reported to outperform state-of-the-art (SOTA) single-sequence RNA LMs in diverse downstream tasks, especially attention-based zero-shot RNA secondary structure prediction, suggesting the significant importance of evolutionary information captured using deep MSAs ^6,7^.

Protein MSAs generated using the Big Fantastic Database (BFD), comprising 2.5 billion reference protein, metagenomic, and metatranscriptomic sequences, contributed to the SOTA accuracy of AlphaFold2 ^8^. Utilization of metagenomic data aided the extraction of co-evolutionary information from MSAs and enhanced prediction of protein tertiary structures, annotations of protein functions, and enzyme discovery ^9^. DMFold, which uses the iterative MSA generation method DeepMSA2, coupled with huge metagenomic resources, achieved improved performance for predicting protein monomers and multimers compared to other SOTA methods in CASP15 ^10^. To efficiently harness the vast scale of raw sequencing repositories, the LOGAN database channeled the information from SRA (Sequence Read Archive) into contigs and unitigs ^11^, leading to >100-fold compression of SRA data, analysis efficiency, and improved quality of the genomic content ^11^. Harnessing the potential of this database using the full 0.9 petabase version or even the 50% sequence identity clustered version (Logan50) resulted in a greater number of effective homologs (N_eff_) and improved structure predictions (pLDDT scores) of viral protein structures ^11,12^, when combined with ColabFold ^13^. Additionally, several teams in CASP16 leveraged custom MSAs from metagenomic resources and databases to improve MSA depth and facilitate structure predictions ^14,15^.

Recent frameworks demonstrate that expanding sequence database scale and diversity directly influences downstream performance. For RNA 3D structure prediction, NuFold ^16^ utilized MSAs generated using rMSA ^17^ alongside a combination of metagenomic sequences from different resources, including NCBI env_nt, MGnify ^18^, and TARA Ocean Metagenome ^19^, deepen alignments and enhance modeling accuracy. Similarly, foundational RNA language models have scaled up pretraining data; Uni-RNA^20^ achieved SOTA performance across multiple tasks using a proprietary, 1-billion-sequence database modeled after MARS_v1.0 ^4^, while AIDO-RNA utilized the 886-million-sequence MARS50 (a 50% identity clustered version of MARS_v1.0) dataset. Notably, while the noisy nature of MARS50 caused AIDO.RNA-1B^21^ to underperform an RNAcentral-pretrained variant in secondary structure tasks, the use of this database enhanced cross-species splicing site prediction. Nevertheless, these training collections remain fundamentally constrained by legacy database coverage and redundancy profiles, perpetuating a homology bottleneck for orphan RNAs and limiting downstream generalization in advanced 3D structure prediction and RNA design.

Ultimately, because contemporary RNA sequence databases are either outdated or entirely proprietary, a comprehensive, publicly available repository is critically needed to leverage tools like rMSA^17^ and RNAcmap3^4^ for training next-generation RNA LMs and accurate deep-learning structural predictions.

Here, we present an elevated database of RNA sequences called “elDORS_raw”, which not only updates, but also expands, rebalances, and deduplicates its integrated data sources, optimizing the database for the MSA generation pipelines and for next-generation RNA foundation models. With respect to the number of sequences, the elDORS_raw database is ∼19 times bigger than the NCBI nucleotide (nt) database and ∼1.11 times bigger than MARS_v1.0. We benchmarked the database using an 80% identity clustered version (elDORS) comprising sequences of length 10 up to 4096 nucleobases, optimized separately for generating deep MSAs more efficiently with both rMSA and RNAcmap3 methods. elDORS is ∼7 times smaller than elDORS_raw and ∼2.7 and ∼2.5 times smaller than MARS_v1.0 and NCBI nt databases, but with improved average MSA depth and computational efficiency. From benchmarking two state-of-the-art methods for MSA generation (rMSA and RNAcamp3) using elDORS, we demonstrate that elDORS provides deeper MSAs with more diverse homologs compared to those obtained using MARS_v1.0 or NCBI nt. Overall, elDORS_raw and its optimized clustered version elDORS are designed to support more sensitive homolog detection and more informative coevolutionary signals. This will improve secondary and tertiary structure inference, functional annotation, and data efficiency in large-scale RNA representation learning.

## Results

### Building and optimization of elDORS

The elDORS_raw database developed in this work has a disk size of 4047.85 GB, comprising 1918768856 (∼1.91 billion) sequences. elDORS_raw comprises all the available genomic, ncRNA, and metagenomic sequences (**Table 1; Fig.1**). Due to its availability in the multivolume gzipped FASTA format, this database is adaptable for generating an MSA using diverse MSA generation methods.

**Table 1.** List of the databases included in elDORS.

| Database | Source | Download date |
| --- | --- | --- |
| nt (nucleotide) | https://ftp.ncbi.nlm.nih.gov/blast/db/ | June 16, 2025 |
| nt subsets:<br>env_nt_v4<br>env_nt_v5<br>Pat_nt (v5)<br>tsa_nt (v5) | https://ftp.ncbi.nlm.nih.gov/blast/db/v4/<br>https://ftp.ncbi.nlm.nih.gov/blast/db/v5/ | June 14, 2025 |
| RNAcentral | https://ftp.ebi.ac.uk/pub/databases/RNAcentral/current_release/sequences/ | June 16, 2025 |
| Rfam | http://ftp.ebi.ac.uk/pub/databases/Rfam/CURRENT/fasta_files/ | June 13, 2025 |
| MGnify | https://ftp.ebi.ac.uk/pub/databases/metagenomics/mgnify_genomes/ | Between March 6-14, 2025 |
| IMGVR | https://genome.jgi.doe.gov/portal/IMG_VR/IMG_VR.home.html | March 16, 2025 |
| GWH | https://download.cncb.ac.cn/gwh/ | Between March 11-13, 2025 |
| GTDB<br>(refseq, GCF) | https://data.ace.uq.edu.au/public/gtdb/data/releases/release226/ | April 16, 2025 |
| gcMeta | https://gcmeta.wdcm.org/ | March 20, 2025 |
| TARA Ocean<br>Metagenome-<br>reconstructed<br>MAGs | https://figshare.com/articles/dataset/The_Reconstruction_of_2_631_Draft_Metagenome-Assembled_Genomes_from_the_Global_Oceans/5188273/13 | June 16, 2025 |
| Greenegenes2 | https://ftp.microbio.me/greengenes_release/current/ | June 17, 2025 |
| MARS-v1.0 | https://ngdc.cncb.ac.cn/omix/release/OMIX003037 | February 28, 2025 |

**Fig. 1.**
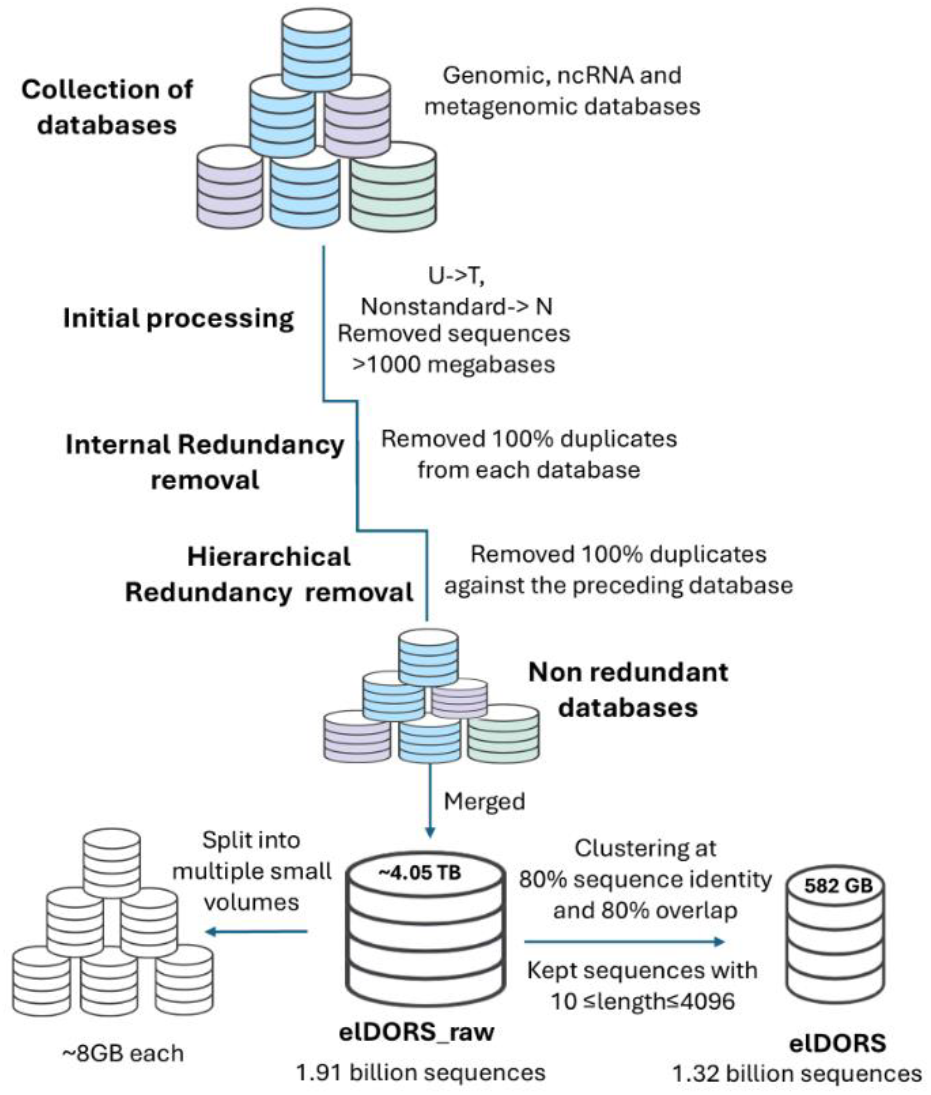
elDORS database preparation workflow schema. Schematic of sequential filtering, internal and hierarchical (redundancy removal against each preceding database listed in **Table 1**) deduplications, and sequence clustering workflows used to compress raw multi-source genomic and metagenomic data (∼4.05 TB, 1.91 billion sequences) into the non-redundant elDORS database (582 GB, 1.32 billion sequences). Specific processing thresholds, sequence length constraints, and volume-splitting parameters are detailed at each progressive stage of database refinement.

Similar to Chikhi and coworkers, who demonstrated that a greater number of effective protein homologs (Neff) could be achieved with a 50% sequence-identity clustered version (Logan50) ^11^, we built and optimized a smaller version of elDORS_raw, called “elDORS”. While the large sequence corpus of elDORS_raw shall be used or further optimized as per the users’ pre-processing preferences required for diverse downstream tasks (e.g., large-scale RNA language model training), elDORS was clustered using 80% sequence identity and 80% coverage thresholds. It has a disk size of 582 GB and comprises ∼1.32 billion non-redundant sequences (10 <= length <= 4096). To rigorously assess how this reduction in database size impacts alignment quality, we evaluated the performance of the elDORS-augmented MSA generation pipelines (indicated throughout by the “-elDORS” suffix to denote database integration) across three datasets (**Fig.2**).

**Fig. 2.**
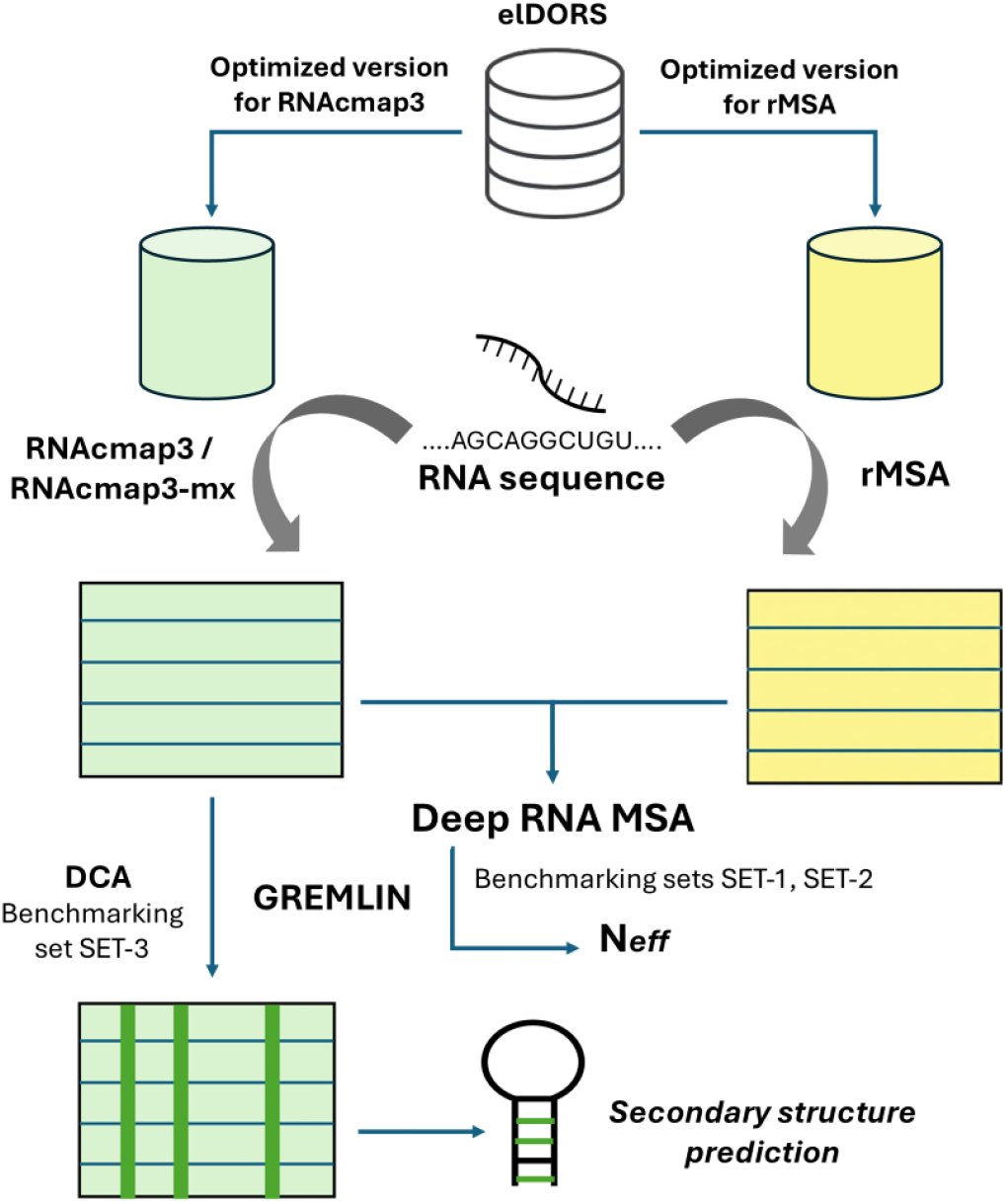
elDORS database benchmarking workflow schema. Workflow illustrating the comparative evaluation of optimized elDORS database variants across distinct downstream prediction modalities. Input target sequences drive multiple sequence alignment (MSA) generation via RNAcmap3, RNAcmap3-mx, or rMSA architectures. Resulting alignments are validated for MSA depth (*Neff*) using benchmarking sets SET-1 and SET-2, or SET-3 to evaluate MSA depth and secondary structure prediction accuracy.

### Effective RNA MSA generation using elDORS-augmented pipelines

Our initial evaluation of elDORS focused on a benchmarking SET-1 dataset, which we specifically curated from 72 PDB sequences (80≤length≤400). SET-1 includes diverse RNAs such as ribosomal fragments, tRNAs, riboswitches, and viral elements, as well as some engineered constructs. The results for SET-1 reveal crucial differences between the two MSA generation methods and their performance when used with the elDORS database. Comparison of the N_eff_ and log_10_(N_eff_) values suggest significantly improved performance of both rMSA and RNAcmap3 approaches with elDORS80 (and RNAcentral as db0 for rMSA) compared with baseline rMSA with the nt (and RNAcentral as db0) database.

Both elDORS-augmented pipelines yielded highly significant improvements over baseline rMSA (paired Wilcoxon signed-rank test; *p*-value = 3.37 ×10^-6^ for rMSA-elDORS, and 3.32 ×10^-8^ for RNAcmap3-elDORS) **(Fig.3A)**. This systemic enhancement across the dataset is also reflected in the summary metrics (**Table S4**), where RNAcmap3-elDORS achieved the highest overall mean log_10_(N_eff_) of 2.879 (median = 2.783) compared to the baseline rMSA (mean = 2.378 and median = 2.364). The rMSA-elDORS variant also demonstrated consistent, albeit intermediate, structural depth gains with an elevated mean of 2.567 and a median of 2.617. For orphan type (i.e., low N_eff_) sequences 1FOQ_A, 2KRL_A, 4P8Z_A, 6WLK_A, and 8T2O_R, for which baseline rMSA showed log_10_(N_eff_) <1, the elDORS-augmented methods (particularly RNAcmap3) were observed to show significantly greater log_10_(N_eff_) values (**Table S4**). Interestingly for 7JRT_A, where rMSA resulted in log_10_(N_eff_)=0 even with elDORS, RNAcmap3-elDORS resulted in log_10_(N_eff_) >3. For 23 sequences, the RNAcmap3-elDORS approach led to the predefined saturation limit of 50,000 sequences. This saturation also underscores an algorithmic distinction between the MSA and RNAcmap3 approaches and their utilization of elDORS. RNAcmap3 employs a broad-sweep heuristic that extracts a maximal initial pool of alignments (num_alignments=50000), which the elDORS database then populates with diverse evolutionary homologs, yielding high effective sequence depths. On the other hand, rMSA employs a multi-stage iterative search governed by predefined sufficiency thresholds (Nf ≥ Nf_cut), yielding comparatively shallow alignments despite querying the same database. This algorithmic distinction is further reflected by the cumulative distribution and per-sequence performance profiles (**Fig.3C, 3E**). Particularly, from the cumulative fraction plot (**Fig.3C**), we observed that while baseline rMSA and rMSA-elDORS leave a notable fraction of sequences below log_10_(N_eff_) = 1.0, the RNAcmap3-elDORS workflow completely shifts the population curve to the right, ensuring that all the sequences in SET-1 exceed this value. The performance comparison for individual sequences (**Fig.3E**) suggests that the specific orphan-type (log_10_(N_eff_) < 1.0) targets, observed with baseline rMSA (highlighted in green), undergo sharp increases in alignment depth when routed through the RNAcmap3-elDORS pipeline.

**Fig. 3.**
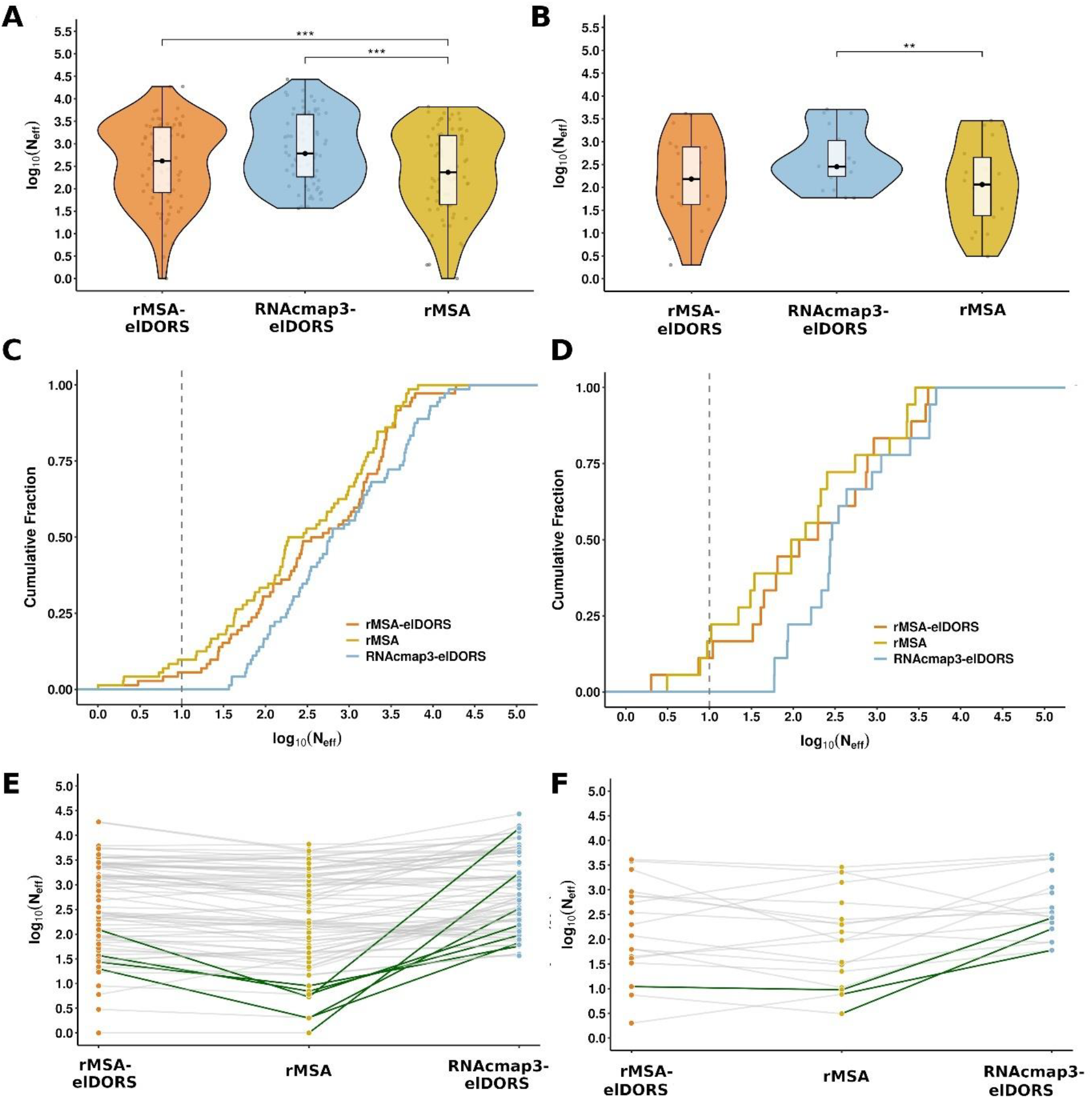
**(A, B)** Violin plots displaying the log_10_(N_eff_) for rMSA-elDORS (orange), RNAcmap3-elDORS (light blue), and baseline rMSA (golden yellow) across benchmarking datasets (**A**) SET-1 (natural RNA sequences) and (**B**) SET-2. Within each violin plot, the box plot indicates the interquartile range, with the solid black dot representing the median, while semi-transparent gray points represent values for individual RNA sequences. The statistical significance between the elDORS-augmented (indicated by the “-elDORS” suffix) pipelines and the baseline rMSA method was determined using a paired Wilcoxon signed-rank test (** p < 0.01; *** p < 0.001). Brackets indicate significant differences; non-significant comparisons are excluded for clarity. **(C, D)** Empirical cumulative distribution function (ECDF) plots showing the cumulative fraction of sequences at varying log_10_(N_eff_) values for (**C**) SET-1 and (**D**) SET-2. The vertical dashed gray line indicates the threshold of log_10_(N_eff_) = 1.0 used to define orphan sequences. **(E, F)** Paired slope graphs centered on the rMSA baseline visualizing independent, sequence-specific performance changes for (**E**) SET-1 and (**F**) SET-2. Dark green lines highlight “rescued” orphan sequences (baseline log_10_(N_eff_) < 1.0 successfully improved to ≥1.0 by the elDORS-augmented pipelines), while semi-transparent gray lines represent all other sequences.

### elDORS-mediated improvement of MSA depth for CASP RNA targets

For the CASP-derived benchmarking SET-2, RNAcmap3 coupled with elDORS generated deeper MSAs, providing a significantly greater number (paired Wilcoxon signed-rank test; *p*-value = 0.0023) of effective sequences **(Fig.3B)**. RNAcmap3-elDORS achieved an overall mean log_10_(N_eff_) of 2.625 and a median of 2.452, greater than the baseline rMSA (mean = 2.052, median = 2.062) (**Table S5**). In contrast, the rMSA approach, when used with elDORS, performed comparably to the baseline without a statistically significant difference (*p*-value = 0.369), resulting in a mean log_10_(N_eff_) of 2.199 and a median of 2.183 (**Table S5**). The overall performance of the elDORS-augmented MSA generation pipelines across both datasets confirms that integrating the elDORS database either substantially enhances or reliably maintains sequence retrieval depth.

For the orphan sequences within this dataset, RNAcmap3-elDORS demonstrated particularly notable improvements in homolog retrieval capacity. For example, the CASP16 target 9FCN_A, baseline rMSA yielded only 17 sequences with a log_10_(N_eff_) of 0.492, whereas the RNAcmap3-elDORS approach expanded the alignment to 177 sequences, achieving a log_10_(N_eff_) of 2.212 (**Table S5**). Similarly, for the target R1293, the baseline rMSA recovered a mere 25 homologs (log_10_(N_eff_) = 0.885). While rMSA-elDORS struggled with this specific target (retrieving only 2 sequences), RNAcmap3-elDORS successfully generated an MSA of 60 sequences with a significantly improved log10(Neff) of 1.778. This observation suggests the suitability of RNAcmap3 for generating MSAs for such orphan-type sequences while utilizing the non-redundant elDORS database architecture.

Finally, RNAcmap3-elDORS successfully reached the 50,000-sequence saturation limit for CASP targets R1241 (Group II intron) and R1271 (RECC component), belonging to deep-homology structural families, highlighting its continued capability to deeply mine the elDORS architecture. The cumulative distribution (**Fig.3D**) highlights that RNAcmap3-elDORS successfully eliminates the low-N_eff_ tail seen in the other two workflows, lifting the entire dataset to log_10_(N_eff_) ≥ 1. These sequence-specific improvements are clearly visualized in the performance comparison for individual sequences (**Fig.3F**), where the green-highlighted lines showcase the substantial improvement achieved using the RNAcmap3-elDORS workflow.

Benchmarking results for SET-1 and SET-2 suggest that the RNAcmap3 approach generally retrieves a much larger number of homologous sequences, hence resulting in a greater effective number of sequences (Neff) than the rMSA method for the same database (elDORS) **(Tables S4, S5, Fig.3)**.

### Direct b*enchmarking performance of RNAcmap3 with elDORS vs MARS*

To pinpoint the performance gain specifically attributable to the elDORS database, we also compared elDORS-augmented pipelines directly against the results for the Rfam-mapped sequences from the RNAcmap3-MARS method and Rfam-derived ^5^ MSAs ^4^.

The performance on SET-3 (**Table S6**) highlights the critical role of the initial 2D structure prediction in guiding the homology search. Here, we evaluated an additional variant, RNAcmap3-mx coupled with elDORS, which utilizes the deep-learning-based MXfold2 ^22^ for 2D structure initialization instead of the standard RNAfold ^23^. While highly accurate standalone deep-learning models like SPOT-RNA2 exist, they are often computationally prohibitive for iterative large-scale database searching and risk overfitting to specific PDB motifs. We selected MXfold2 and RNAfold because these methods provide computationally efficient, thermodynamically grounded structural “seeds” that generalize robustly across diverse evolutionary landscapes. Although MXfold2’s training set includes PDB-derived data, which could introduce potential data leakage, its inclusion here serves to determine if a more accurate initial structural profile further enhances the retrieval of distant homologs from elDORS.

For SET-3, RNAcmap3-elDORS resulted in a mean log_10_(N_eff_) of 2.485 (median = 2.441), performing comparably to RNAcmap3-MARS (mean = 2.504, median = 2.396) without a statistically significant difference (paired Wilcoxon signed-rank test; p-value = 0.932). Both pipelines, however, yielded a significantly greater MSA depth than Rfam (mean = 1.473, median = 1.472). The RNAcmap3-mx-elDORS approach achieved the highest overall mean log10(Neff) of 2.631 and a median value of 2.669, though this also remained statistically comparable to the MARS baseline (p-value = 0.149) (**Fig.4A, Table S7A**). Similar to the observation for RNAcmap3 coupled with the MARS database, the RNAcmap3 and RNAcmap3-mx pipelines coupled with elDORS exhibited a higher minimum sequence yield across all tested RNA sequences in SET-3. Hence, the direct comparison revealed the effective elimination of the ‘low-yield’ (poor N_eff_) failures frequently observed earlier in Rfam and RNAcmap2 ^4^.

**Fig. 4.**
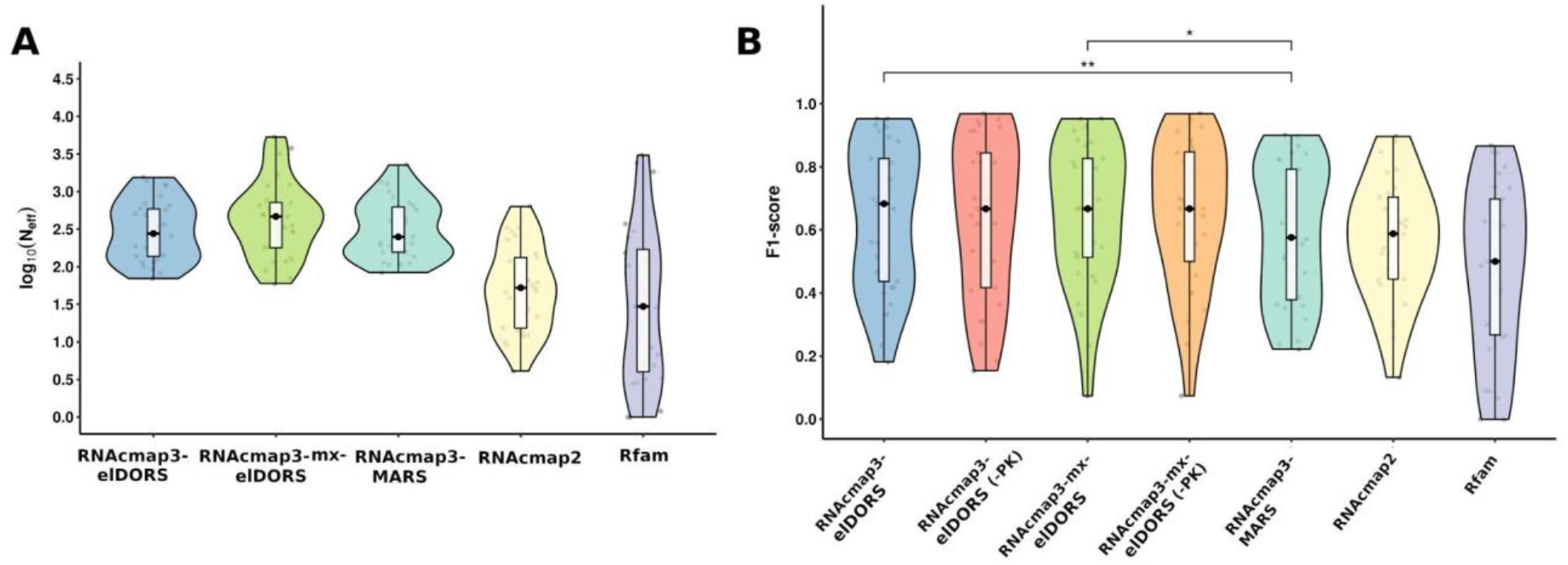
(**A**) Violin plots of log_10_(N_eff_) for the benchmarking dataset SET-3. The elDORS-augmented pipelines perform comparably with RNAcmap3, coupled with MARS. (**B**) Violin plots of F1-scores for RNA secondary structure prediction for the benchmarking dataset SET-3. Statistical significance between the results from elDORS-augmented (indicated by the “-elDORS” suffix) pipelines (considering pseudoknots) and RNAcmap3-MARS^4^ was determined using a paired Wilcoxon signed-rank test (* p < 0.05, ** p < 0.01). Within each violin plot, the box plot indicates the interquartile range with the solid black dot representing the median, while semi-transparent gray points represent values for individual RNA sequences. In panel B, “(-PK)” indicates calculation of F1-score without considering pseudoknots.

RNAcmap3-elDORS achieved a mean F1-score of 0.638 (median = 0.683), representing a highly significant improvement over the RNAcmap3-MARS baseline (paired Wilcoxon signed-rank test; p-value = 0.0014). This performance also significantly outpaced RNAcmap2 (mean = 0.567; median = 0.588) and Rfam (mean = 0.472; median = 0.500) (**Fig.4B, Table S7A**). The 2D structural initialization via MXfold2 (RNAcmap3-mx-elDORS) pushed the mean F1-score even higher to 0.647 (median = 0.667), demonstrating a better performance compared to the RNAcmap3-MARS approach (p-value = 0.0345). Interestingly, focusing exclusively on general base pairs by omitting pseudoknot predictions (-PK) maintained comparable overall F1-scores (**Fig.4B, Table S7A**).

Because we utilized the standard threshold of extracting the top L/3 predicted pairs from direct-coupling analysis (DCA), Positive Predictive Value (PPV) and F1-scores are inherently deflated for sequences where the true number of base pairs is less than the L/3 cutoff. Consequently, Sensitivity (Recall) serves as the more crucial and direct metric for evaluating a pipeline’s ability to successfully recover evolutionary base-pairing signals. For the RNAcmap3-mx-elDORS pipeline, evaluation without considering PKs increased mean sensitivity (recall) from 0.693 to 0.759, while mean PPV (precision) correspondingly decreased from 0.634 to 0.607 (**Fig.5A, B, Table S7B**). This suggests that while filtering out pseudoknot topologies slightly reduces precision, it allows the model to capture a substantially broader set of true canonical interactions for the sequences in SET-3. To visualize the co-dependency of these metrics, we employed a parity-anchored scatter plot (**Fig.5C**), which reveals that the values for the majority of the SET-3 sequences reside in the high-performance region. This observation highlights that despite the conservative L/3 prediction threshold, the elDORS-augmented pipelines frequently achieve near-maximal sensitivity while remaining proximal to the parity line, demonstrating the reliability of the structural signals extracted by elDORS.

**Fig. 5.**
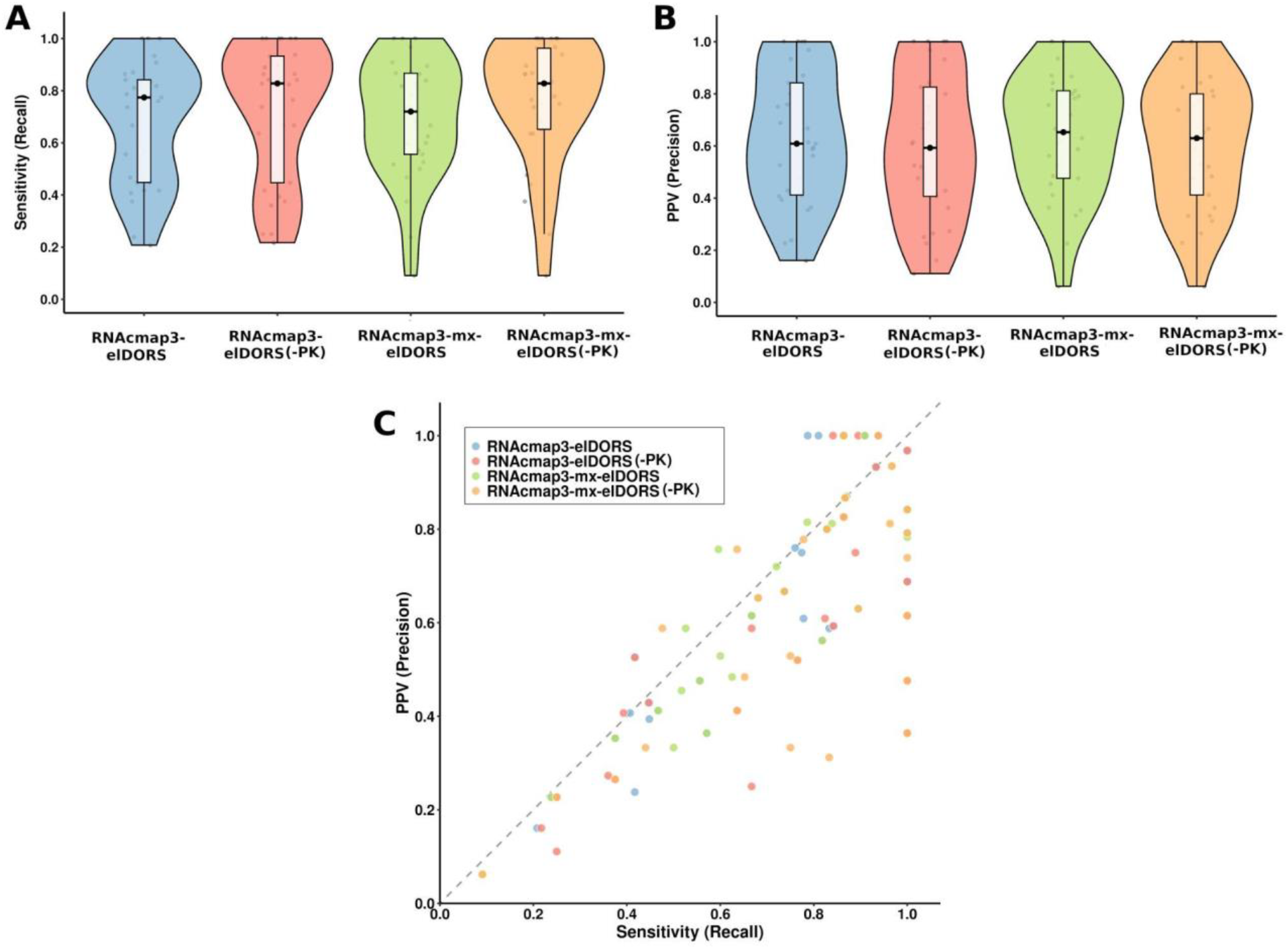
Violin plots (central box plots indicate interquartile range with the solid black dot representing the median and semi-transparent gray points representing values for individual RNA sequences) of (**A**) Sensitivity (Recall) and (**B**) PPV (Precision) for the benchmarking dataset SET-3. (**C**) Scatter plot illustrating the co-dependency between Sensitivity and PPV for each sequence, grouped by method. The elDORS-augmented pipelines are indicated by the “-elDORS” suffix and “(-PK)” indicates calculation of metrics without considering pseudoknots.

## Discussion

The recent revolution in protein structure prediction demonstrated that the depth and diversity of underlying evolutionary data are crucial for computational predictive accuracy. The RNA field has long been constrained by fragmented repositories and the homology bottleneck of orphan sequences. We designed elDORS_raw to unify metagenomic and transcriptomic data along with diverse existing genomic and non-coding RNA databases into a single, high-performance database.

elDORS, our lightweight yet information-dense 80% identity clustered version of elDORS_raw, results from internal redundancy removal, hierarchical redundancy removal, filtering, and clustering to ensure its seamless multipurpose usage. In that sense, elDORS supersedes the MARS_v1.0 database (1.57 TB). Because we optimized and preprocessed elDORS for its utilization with RNAcmap3 and rMSA, users will more efficiently generate RNA MSAs. In a nutshell, elDORS enables more sensitive homology detection and more accurate structural predictions, despite being significantly smaller than MARS or NCBI nt. Altogether, elDORS is sufficient to eliminate the shallow MSA depth issues that often hinder automated MSA-based approaches. Using elDORS, we demonstrate that it is possible to achieve deeper MSAs and higher structural F1-scores while reducing computational overhead.

The landscape of automated RNA multiple sequence alignment (MSA) pipelines has evolved significantly to balance computational efficiency with the retrieval of deep, informative homologies. Early approaches, such as the original RNAcmap ^24^ pipeline, established a baseline by pairing BLAST-N ^25^ with Infernal ^26^ covariance models (CM) against standard NCBI databases to extract evolutionary couplings for structure prediction tools like SPOT-RNA2 ^27^. To capture richer evolutionary signals, subsequent methods expanded both their search depth and breadth. For instance, RNAcmap2^28^ introduced iterative Infernal searches and incorporated diverse environmental and transcriptome databases, while the rMSA ^17^ method adopted an exhaustive five-iteration search across Rfam, RNAcentral, and the NCBI nt database. Recognizing the need to consolidate these massive, fragmented sequence pools into a more manageable framework led to the development of the MARS database more recently ^4^. By coupling this structurally focused database with the updated RNAcmap3 algorithm (RNAcmap3-MARS), Chen and coworkers reported a more robust and targeted homology search workflow.

Our benchmarking results suggest that computational efficiency and predictive accuracy are not solely the result of database scale, but also of curation quality and non-redundancy. The superior performance of RNAcmap3 (or its recommended MXfold2-assisted variant RNAcmap3-mx) and rMSA when coupled with elDORS indicate that redundant sequence space often behaves as an evolutionary noise that can trigger early-stopping thresholds in iterative search heuristics.

elDORS also serves as the foundational infrastructure for the next generation of RNA foundation models and deep-learning-based RNA design tools. As large-scale RNA LMs increasingly move toward MSA-based architectures to capture long-range dependencies, the availability of a robust, up-to-date, and non-redundant resource like elDORS is essential for pre-training and zero-shot structural inference.

Through building this robust, ready-to-use sequence database and clearly outlining the data collection, processing, and optimization methodology, our study aims to facilitate further efficient downstream applications of this resource. Crucially, by making both the optimized elDORS and the complete elDORS_raw database freely available, we aim to accelerate the pace of discovery in RNA biology, providing the community with a versatile toolkit to explore the vast, unannotated diversity of the RNA world and translate evolutionary information into functional insight.

## Methods

### Database preparation

Our method of creating elDORS_raw builds on and improves prior efforts by Chen et al. (2024) ^4^, who created the MARS genomic database, integrating well-annotated nucleotide sequences as well as poorly understood metagenomic sequences. elDORS_raw represents an extensive, up-to-date database, along with more robust and well-optimized versions for efficiency in diverse downstream applications.

### Data collection workflow

For assembling elDORS_raw, we first obtained the databases listed in **Table 1**. We collected genomic sequences from the NCBI nt database and its subsets (env_nt, pat_nt, and tsa_nt) ^1^. Non-coding RNA sequences from the RNAcentral database^2,3^ and a comprehensive, curated set of sequences representing the known RNA families, assembled in the Rfam ^5^, were also collected. Bacterial and Prokaryotic genomes were collected from the GTDB database ^29^, specifically restricting the dataset to highly curated RefSeq (GCF) assemblies. Additionally, 16S rRNA and corresponding genome sequences were sourced from Greengenes2 ^30^. We obtained metagenomic sequences from MGnify ^31^ and the Global Catalogue of Metagenomics (gcMeta) ^32^ databases. Viral sequences (corresponding to viral genomes, metagenomes, and metatranscriptomes) were collected from the IMG/VR database ^33^, and additional genomic sequences were sourced from the Genome Warehouse (GWH) ^34^. Marine sequences were retrieved from a published dataset of 2,631 draft metagenome-assembled genomes reconstructed from the TARA Oceans expedition ^19^. The individual sequences were assembled (merged into a single FASTA file) using in-house Python scripts, where needed for a specific database.

Due to the issues with the authentication server of MG-RAST, we were unable to fetch the transcriptomic and metagenomic sequences directly from the corresponding portal. It’s also important to note that due to such issues, it remains challenging to perform regular/periodic updates of the database without meticulous human intervention and time. Hence, we extracted the sequences that are present in the original MARS v1.0 database (considering the inclusion of MG-RAST sequences available in MARS v1.0) and not in the component databases that we considered. Additionally, we ensured that we integrated the latest available versions of Greengenes2 (which is included in MG-RAST) separately.

### Data Processing Workflow

The NCBI nt database and the nt subsets were initially retrieved in the NCBI-BLAST format. Subsequently, the FASTA format files corresponding to the respective databases were extracted using the blastcmd command available in the ncbi-blast-2.16.0+ software package ^35^. The RNAcentral database was downloaded in the zipped FASTA format and extracted to obtain the FASTA format file. The nt and RNAcentral databases were processed using the fastaNA code, similar to the protocol followed for the rMSA database curation by Zhang et al. (2023) ^17^. Sequences longer than 1000 megabases were removed, and further processing was performed for each database. First, U residues (if present) were replaced with T residues, followed by the replacement of any non-standard residues (i.e., other than A/T/G/C), including dashes and gaps, with ‘N’. We utilized seqKit ^36^ to remove duplicate (100%) sequences (i.e., internal redundancy removal) from most individual databases, namely Rfam, RNAcentral, SILVA, GTDB, Geengenes2, IMG-VR, gcMeta, TARA ocean metagenome, Mgnify, NCBI tsa_nt, and NCBI patnt. However, due to memory issues faced with the seqKit tool for env_nt_v4, GWH, and NCBI nt, we employed an in-house Python code for the removal of 100% duplicated sequences from these three databases. For verification that the results obtained from seqKit and the in-house code are the same, we performed an initial test for the Rfam database before applying it to the larger databases.

Next, we followed the same hierarchical approach as was followed by Chen et al (2024) ^4^ to maintain data provenance and performed the removal of 100% duplicated sequences (i.e., hierarchical redundancy removal) from a database in **Table 1** against the database/databases that are on top of it. To account for the sequences (especially from MG-RAST, which were present in MARS v1.0), each of the 150 volumes from MARS v1.0 was also subjected to redundancy removal against the databases that are listed on top of it in the table. The resulting FASTA files were merged into a single file. However, at this step, we noticed that the resulting database contained “U” and other non-A/T/G/C/N characters, indicating the original MARSv1.0 database contained non-standard characters. Hence, we further processed this merged database, replacing the U residues with T and any non-standard residues (i.e., other than A/T/G/C), including dashes, gaps, and other characters, with ‘N,’ and then subjected the processed database to an additional step of internal redundancy removal. Finally, the resulting databases, as reported in **Table S1**, were merged to create the elDORS_raw database, having a total size of 4047.85 GB containing 1918768856 (∼1.91 billion) sequences, which were split into multiple volumes of gzipped FASTA having a size of ∼8 GB each.

Next, sequences exceeding 4,096 nucleobases or having fewer than 10 nucleobases were filtered out from the resulting component databases (before merging into elDORS_raw). The details of the resulting databases from this filtering step are provided in **Table S2**. Following this, the resulting databases were subjected to clustering using the MMseqs2 linclust algorithm ^37^, where the sequences were clustered at an 80% sequence identity threshold and 80% overlap with the longest sequence in each of the clusters. Then, to combine the databases, we aggregated the individual clustered databases into a master database and performed a final round of MMseqs2 linclust at 80% identity and overlap thresholds. This final step was conducted to eliminate cross-source redundancy, producing a unified set of non-redundant sequences and forming the elDORS80 database, which contains 1323935871 (i.e., ∼1.32 billion) sequences (**Table S3**). Moreover, we noticed that a number of the additional sequences retrieved from the MARS database contained long redundant header sections. To ensure compatibility with the rMSA pipeline, the elDORS database was pre-processed to standardize sequence identifiers. Specifically, complex MG-RAST headers present in the original MARS database were replaced with simplified, unique alphanumeric tags (e.g., _seq1, _seq2) after the first 25 characters from the original identifier, preventing buffer overflow errors and identifier truncation during the BLASTN and Infernal search phases. Users can conduct the identifier processing step when using the elDORS_raw database with rMSA; however, for RNAcmap3, this step can be skipped. To make the elDORS80 database compatible with RNAcmap3, we split the database into 114 volumes, each ∼5GB in size, and converted the merged database to BLAST format per RNAcmap3 guidelines.

### Benchmarking workflow

Following the optimization of elDORS, we benchmarked the utility of this database in three steps (**Fig.2**) to progressively validate the database. In the first step, we curated a dataset ‘SET-1’ comprising 72 sequences (80≤length≤400) from the PDB to benchmark the elDORS database and the methods for generating MSAs. MSAs for these sequences corresponding to the standard rMSA (with the nt and RNAcentral databases) were obtained from the Kaggle Ribonanza part-1 datasets^38^. The selection of the length filter spanning a structurally complex and computationally tractable length regime ensured an optimal benchmarking dataset with sufficient sequence complexity to avoid spurious homology matches in a massive database. RNAcmap3 (with elDORS) and rMSA (with RNAcentral as db1 and elDORS as db2) were carried out with default settings. To effectively compare the rMSA and RNAcmap3 approaches, five synthetic RNAs were additionally included in the assessment, and sequences with crystallization scaffolds were discarded.

In step 2, we considered a dataset ‘SET-2’ comprising 18 RNA target sequences of diverse lengths, from CASP competitions (2 from CASP15 and 16 from CASP16) to evaluate generalization on blind target sequences. Here, MSAs with standard rMSA (with RNAcentral as db1 and nt as db2 databases) were generated using the unprocessed versions of the nt and RNAcentral databases, which were used for preparing elDORS.

In step 3, direct benchmarking was performed considering the 29 Rfam-mapped sequences ‘SET-3’ previously reported for RNAcmap3/MARS benchmarking by Chen et al, 2024 ^4^. Notably, when we investigated the 3D structures of SET-3 sequences, we noticed that some of the structures had missing/unresolved residues. 2IL9_A (present among the 30 sequences tested by Chen et al, 2024 ^4^, also present in our benchmarking dataset SET-1) was discarded from the benchmark in SET-3 due to 7 unresolved nucleotides in the 3D structure. Other sequences with a few missing nucleotides were manually approximated based on their relative positions, and the original 3D-derived 2D structures are shown in **Table S6**. Here, we introduced RNAcmap3-mx (which uses MXfold2 ^22^ for the RNA 2D structure prediction instead of RNAfold ^23^) and performed a thorough comparison of the number of effective sequences (N_eff_) as well as secondary structure signal derivation from Direct Coupling Analysis (DCA). The N_eff_ calculation and DCA were carried out using GREMLIN (https://github.com/sokrypton/GREMLIN_CPP)^39^, considering a threshold value of 0.8 and a gap cutoff of 0.5 for all the alignments generated in this work (following the RNAcmap3 script). The ground truth secondary structures were obtained using the RNApolis ^40^ tool within the RNAPDBee ^41^ webserver or using the RNApolis ^40^ Python program locally for structures with interacting chains. The ground truth secondary structures considered for the F1-score calculation were plotted using VARNA ^42^ (**Table S7**). F1-scores were calculated using an in-house Python script considering the Top L/3 pairing signals. Statistical comparisons of F1-scores and log_10_(N_eff_) between different methods were performed using a two-sided, paired Wilcoxon signed-rank test. Significance thresholds were defined as *p* < 0.05 (*) and *p* < 0.01 (**). All statistical tests and visualizations were generated using R (ggplot2 and ggsignif packages).

## Data Availability

The version-1 of the elDORS database (elDORS_v1) in a multi-volume zipped version (.fasta.gz format) (∼170GB) is available for download at https://gist.github.com/duttan710/7e8d2a251e37ab405eeaac10dd55ccf3#file-eldors_v1_index-md.

## Supporting information

Supplementary material is available online.

## Acknowledgements

We wish to thank Gabriel Galvez for his assistance in managing our GPU workstation; John Gillet and the University of Houston IT department for hosting our GPU workstation; the Research Computing Data Core at the University of Houston for making it possible for us to use the Carya Cluster; AWS and especially Bill Short and Bineesh Ravindran for discussions and support for making elDORS available to the community.

## Funding

This research was supported by start-up funds from the University of Houston (to Q.V.).

## Author contributions

N.D. and Q.V. conceived the study, analyzed performance, interpreted the results, and contributed to the drafting, editing, and final approval of the manuscript. N.D. developed the methodology, implemented the workflow, constructed the database and datasets, and carried out benchmarking.

## Notes

https://gist.github.com/duttan710/7e8d2a251e37ab405eeaac10dd55ccf3#file-eldors_v1_index-md

### Competing Interest Statement

The authors have declared no competing interest.

