## Supplementary material for "elDORS: An elevated Database Of RNA Sequences": SI

**Table S1.** List of the integrated databases and the relevant statistical details for elDORS_raw

| Database | Size in fasta | No. of sequences | No. of nucleotide bases | Max and Min sequence lengths | Avg. sequence length | Final size (GB) (after removal of redundancy) |
|---|---|---|---|---|---|---|
| **db1**<br>NCBI nt* | 1443 GB<br>* 490 GB | 102960590<br>**99944650** | 1528186360278<br>**1521659938132** | 99984324<br>6<br>**99984324**<br>**6** | 14842.44<br>**15225.03** | **1437 GB†**<br>*490 GB |
| **db2**<br>NCBI env_nt_v5<br>env_nt_v4 (last updated 2020.02.03 | 0.85 GB<br>142 GB<br>*113 GB | 470514<br>**470101**<br>124601035<br>**121283291** | 833744959<br>**833470165**<br>134952479671<br>**132219696026** | 8819083<br>7<br>**8819083**<br>**7**<br>40002466<br>6<br>**40002466**<br>**6** | 1771.99<br>**1772.96**<br>1083.08<br>**1090.17** | 0.85 GB†<br>**0.85 GB††**<br>140 GB†<br>**139 GB††**<br>*108 GB |
| **db3**<br>NCBI tsa_nt | 15 GB<br>*15 GB | 16443012<br>**16415243** | 13791718074<br>**13758730107** | 65905<br>14<br>**65905**<br>**14** | 838.76<br>**838.17** | 15 GB †<br>**15 GB††**<br>*14 GB |
| **db4**<br>NCBI pat_nt | 31 GB | 49431229<br>**24618418** | 28481566926<br>**13294837117** | 38042092<br>6<br>**38042092**<br>**6** | 576.19<br>**540.04** | 16 GB †<br>**15 GB††** |

| | | | | | | |
|---|---|---|---|---|---|---|
| | *28 GB | | | | | *13 GB |
| **db5**<br>RNAcentral* | 26 GB<br><br>*18 GB | 32429977<br><br>**22225575** | 26478795555<br><br>**19259631976** | 985945<br>10<br>**985945**<br>**10** | 816.49<br><br>**866.55** | 26 GB †<br><br>**19GB ††** |
| **db6**<br>Rfam | 2.4 GB | 10025911<br><br>**4238848** | 1217268433<br><br>**562052218** | 10799<br>19<br>**10799**<br>**19** | 121.41<br><br>**132.60** | 1.6 GB †<br><br>**1.1 GB††** |
| **db7**<br>MGnify* | 744 GB<br><br>*35GB | 64504541<br><br>**60857656** | 784394318511<br><br>**728421143146** | 7084828<br>51<br>**5631190**<br>**51** | 12160.30<br><br>**11969.26** | 698 GB†<br><br>**688 GB††** |
| **db8**<br>IMG-VR | 153 GB | 15722824<br><br>**15314931** | 160201446086<br><br>**153640862013** | 2473870<br>165<br>**956726**<br>**282** | 10189.10<br><br>**10032.10** | 148 GB†<br><br>**146 GB††** |
| **db9**<br>gcMeta | 95 GB | 12235059<br><br>**12099213** | 99855179622<br><br>**98531213632** | 2619196<br>1000<br>**2619196**<br>**1000** | 8161.40<br><br>**8143.61** | 94 GB †<br><br>**94 GB††** |
| **db10**<br>GWH* | 664 GB<br>After removing >1000Mb sequences (629GB)<br>*387 GB | 32963406<br><br>**31402262** | 666252855154<br><br>**633417721618** | 882488833<br>84<br>**882488833**<br>**84** | 20211.89<br><br>**20171.09** | 601 GB †<br><br>**599GB ††**<br><br>*365 GB |
| **db11**<br>GTDB | 154 GB | 2465254<br><br>**2394168** | 162201553726<br><br>**127068028038** | 14782125<br>20<br>**13758192**<br>**20** | 65795.07<br><br>**53073.98** | 154 GB†<br><br>**121GB††** |
| **db12** | 5.5 GB | 214181 | 5900330323 | 2698294 | 27548.34 | 5.6 GB† |

| | | | | | | |
|---|---|---|---|---|---|---|
| TARA Ocean Metagenome | | **208345** | **5766509769** | 4000<br>**2698294**<br>**4000** | **27677.70** | **5.5 GB††** |
| **db13** Greengenes2 | 3.4 GB | 23467470<br><br>**23423863** | 3331901152<br><br>**3270719299** | 4563<br>90<br>**4563**<br>**90** | 141.98<br><br>**139.63** | 3.4 GB†<br><br>**3.4 GB††** |
| MARS (MARS000-149 volumes)<br><br>**db14** MARS_additional | *1571 GB | *1727789860<br><br>**1483872292** | *1592396862523<br><br>**766691711763** | <br><br>**1000000001** | <br><br>**516.68** | <br><br>**764GB †††** |
| Total size | NA | 1918768856 | | | | 1437+2610.85 =4047.85GB |

Note:

The databases already present in the MARS database (v1.0) are indicated with asterisks, and the sizes reported in Chen et al. (2024) are also noted. *Values obtained from Chen et al. (2024).

†Internal redundancy removed

††Redundancy removed against the databases on top of it

††† Internal redundancy was removed from the merged (and processed to contain ATGCN) MARS_additional database after (hierarchical) redundancy was removed for the individual volumes against the databases listed above in the table. The

The statistics for each database are in columns 3-6: original database without internal redundancy removal (top) and after internal redundancy removal and redundancy removal against the databases on top of it (**bottom, bold**).

### Table S2. Statistical details of the databases after the filtering step.

| Database | No. of sequences | No. of nucleotide bases | Max and Min sequence lengths | Avg. sequence length | Final size (GB) (after filtering) |
|---|---|---|---|---|---|
| **db1**<br><br>NCBI nt* | 79209477 | 110219347148 | 4096<br>10 | 1391.49 | 106 GB† |
| **db2**<br><br>NCBI env_nt_v5 | 461301 | 273194297 | 4096<br>15 | 592.23 | 0.31 GB† |

| | | | | | |
|---|---|---|---|---|---|
| env_nt_v4 (last updated 2020.02.03 | 118095004 | 88654206684 | 4096<br>16 | 750.70 | 98 GB † |
| **db3**<br>NCBI tsa_nt | 16050821 | 11590665714 | 4096<br>14 | 722.12 | 13 GB† |
| **db4**<br>NCBI pat_nt | 24262603 | 6174671803 | 4096<br>10 | 254.49 | 8.2 GB† |
| **db5**<br>RNAcentral* | 21902455 | 12584609046 | 4096<br>10 | 574.58 | 13 GB† |
| **db6**<br>Rfam | 4237382 | 553879946 | 4095<br>19 | 130.71 | 1.1 GB† |
| **db7**<br>MGnify* | 26462910 | 67054612511 | 4096<br>51 | 2533.91 | 64 GB† |
| **db8**<br>IMG-VR | 1451251 | 4354482951 | 4096<br>282 | 3000.50 | 4.3 GB† |
| **db9**<br>gcMeta | 6349258 | 15136600534 | 4096<br>1000 | 2384.00 | 15 GB† |
| **db10**<br>GWH* | 25421927 | 35409902734 | 4096<br>84 | 1392.89 | 35 GB† |
| **db11**<br>GTDB | 858223 | 1273608398 | 4096<br>20 | 1484.01 | 1.3 GB† |
| **db12**<br>TARA Ocean Metagenome | 184 | 744448 | 4000<br>4096 | 4045.91 | 0.000744 GB† |
| **db13**<br>Greengenes2 | 23423860 | 3270705998 | 3771<br>90 | 139.63 | 3.4 GB† |
| **db14**<br>MARS_additional | 1466142713 | 455615430423 | 4096<br>10 | 310.76 | 469 GB† |
| Total size | 1814329369 | 812166662635 | | | 831.6107 GB |

†After the removal of sequences exceeding 4,096 nucleobases or having fewer than 10 nucleobases

The number of nucleobases in each sequence in the original DB (without redundancy removal) is added as a data file in the supplementary files.

**Table S3. Statistical details of the databases after the clustering step.**

| Database | No. of sequences | No. of nucleotide bases | Max and Min sequence lengths | Avg. sequence length | Final size (GB) (after clustering) |
|---|---|---|---|---|---|
| **db1**<br><br>NCBI nt* | 41200664 | 52459776818 | 4096<br>10 | 1273.28 | 50 GB† |
| **db2**<br><br>NCBI env_nt_v5<br><br>env_nt_v4 (last updated 2020.02.03 | <br><br>394511<br><br>97074805 | <br><br>239975854<br><br>71953876960 | <br><br>4096<br>15<br><br>4096<br>16 | <br><br>608.29<br><br>741.22 | <br><br>0.268 GB†<br><br>79 GB† |
| **db3**<br>NCBI tsa_nt | 13759476 | 8961418365 | 4096<br>14 | 651.29 | 9.4 GB† |
| **db4**<br>NCBI pat_nt | 17869232 | 4131844727 | 4096<br>10 | 231.23 | 5.6 GB† |
| **db5**<br>RNAcentral* | 7665977 | 4645501519 | 4096<br>10 | 605.99 | 4.5 GB† |
| **db6**<br>Rfam | 928142 | 158462051 | 4095<br>19 | 170.73 | 0.265 GB† |
| **db7**<br>MGnify* | 18860882 | 43832436727 | 4096<br>51 | 2323.99 | 42 GB† |
| **db8**<br>IMG-VR | 1081019 | 3319864429 | 4096<br>282 | 3071.05 | 3.2 GB† |
| **db9**<br>gcMeta | 5770140 | 13598978708 | 4096<br>1000 | 2356.78 | 13 GB† |
| **db10**<br>GWH* | 22458452 | 31235240844 | 4096<br>84 | 1390.80 | 31 GB† |
| **db11**<br>GTDB | 787026 | 1181870877 | 4096<br>20 | 1501.69 | 1.2 GB† |
| **db12**<br>TARA Ocean Metagenome | 184 | 744448 | 4096<br>4000 | 4045.91 | 0.000733 GB† |
| **db13**<br>Greengenes2 | 10479088 | 1206320244 | 3771<br>90 | 115.12 | 1.3 GB† |
| **db14**<br>MARS_additional | 1168968724 | 379469444543 | 4096<br>10 | 324.62 | 386 GB† |

| | | | | | |
|---|---|---|---|---|---|
| Total<br><br>elDORS | 1323715880 | 574498600231 | 4096<br>10 | 434.00 | 588.9338 GB ††<br><br>582 GB* (575 GB**)<br><br>170GB (zipped)<br><br>441 GB (36 volumes) |

†After clustering sequences with at least 80% identity and 80% overlap with the longest sequence in each cluster, resulting in the DB80 versions.

††Sum of the sizes of the individual clustered databases
* Final disk size of the merged and clustered database
** After FASTA sequence header processing

**Table S4. Comparison of the depths of the MSAs generated for the benchmarking dataset SET-1.**

| | **rMSA (with RNAcentral+ elDORS)**<br>**RNAcmap3 (+ elDORS)** | | | Standard rMSA (RNAcentral+nt) | | |
|---|---|---|---|---|---|---|
| Sequence_ID_Chain_ID Length | Number of sequences | Neff | $log_{10}$(Neff) | Number of homologs | Neff | $log_{10}$(Neff) |
| 1FFK_9<br>122 | **8811**<br>**50000** | **1493.64**<br>**4010.87** | **3.174**<br>**3.603** | 7500 | 1585.96 | 3.200 |
| 1FOQ_A<br>120 | **48**<br>**92** | **27.277**<br>**57.164** | **1.435**<br>**1.757** | 13 | 9 | 0.954 |
| 1S9S_A<br>101 | **168**<br>**559** | **38.7449**<br>**338.32** | **1.588**<br>**2.529** | 345 | 42.8848 | 1.632 |
| 1WZ2_D<br>88 | **25127**<br>**50000** | **3630.28**<br>**2920.91** | **3.562**<br>**3.465** | 25191 | 3615.97 | 3.558 |
| 2GO5_9<br>90 | **5598**<br>**34228** | **1048.55**<br>**1313.17** | **3.020**<br>**3.118** | 5378 | 968.368 | 2.986 |
| 1YSH_B<br>101 | **5269**<br>**50000** | **123.47**<br>**630.723** | **2.091**<br>**2.799** | 6232 | 180.358 | 2.256 |
| 2IL9_A<br>142 | **493**<br>**368** | **273.187**<br>**208.401** | **2.436**<br>**2.319** | 161 | 73.1934 | 1.864 |
| 2J28_A<br>117 | **12165**<br>**45720** | **2766.99**<br>**2651** | **3.442**<br>**3.423** | 7700 | 1667.49 | 3.222 |
| 2J37_A<br>128 | **7208**<br>**15579** | **2420.57**<br>**4561.89** | **3.384**<br>**3.659** | 6336 | 2180.2 | 3.338 |

| | | | | | | |
|---|---|---|---|---|---|---|
| 2NBX_A<br>108 | **230**<br>**241** | **68.0707**<br>**112.992** | **1.833**<br>**2.053** | 745 | 75.4003 | 1.877 |
| 2KRL_A<br>102 | **8**<br>**76** | **3**<br>**67.333** | **0.477**<br>**1.828** | 8 | 2.047 | 0.311 |
| 2N1Q_A<br>155 | **2900**<br>**2692** | **202.524**<br>**276.576** | **2.306**<br>**2.442** | 4977 | 404.41 | 2.607 |
| 2NOQ_A<br>190 | **859**<br>**855** | **573.344**<br>**548.31** | **2.758**<br>**2.739** | 341 | 160.821 | 2.206 |
| 2OM7_C<br>96 | **8081**<br>**50000** | **1139.96**<br>**1630.68** | **3.056**<br>**3.212** | 5823 | 659.831 | 2.819 |
| 2XXA_F<br>106 | **12296**<br>**3578** | **5601.12**<br>**1181.15** | **3.748**<br>**3.072** | 6849 | 3528.49 | 3.548 |
| 3IYR_A<br>349 | **27378**<br>**50000** | **18400.9**<br>**4738.23** | **4.264**<br>**3.675** | 6131 | 4762.07 | 3.678 |
| 3NDB_M<br>136 | **4891**<br>**36347** | **2756.14**<br>**9049.23** | **3.440**<br>**3.957** | 6557 | 3530.13 | 3.548 |
| 3IZ4_A<br>377 | **24583**<br>**50000** | **18765**<br>**11323.3** | **4.273**<br>**4.054** | 6501 | 5114.01 | 3.708 |
| 3J3V_B<br>119 | **10867**<br>**50000** | **1659.47**<br>**6039.81** | **3.219**<br>**3.781** | 6107 | 920.748 | 2.964 |
| 3W1K_F<br>101 | **16608**<br>**50000** | **2553.46**<br>**9266.93** | **3.407**<br>**3.967** | 19893 | 3617.86 | 3.558 |
| 4C4Q_N<br>233 | **1638**<br>**1960** | **82.9808**<br>**251.787** | **1.919**<br>**2.401** | 4605 | 130.908 | 2.117 |
| 4P8Z_A<br>188 | **31**<br>**342** | **19.8679**<br>**330.333** | **1.298**<br>**2.519** | 4 | 2 | 0.301 |
| 4V6W_A7<br>120 | **31219**<br>**50000** | **1610.85**<br>**15516.4** | **3.207**<br>**4.191** | 22601 | 535.861 | 2.729 |
| 4TZP_C<br>102 | **2696**<br>**524** | **2157.7**<br>**413.075** | **3.334**<br>**2.616** | 743 | 542.654 | 2.734 |
| 5MC6_BR<br>121 | **34660**<br>**50000** | **2622.98**<br>**8935.62** | **3.418**<br>**3.951** | 18466 | 441.848 | 2.645 |
| 5Z3G_B<br>158 | **4284**<br>**50000** | **849.431**<br>**1455.3** | **2.929**<br>**3.163** | 4159 | 1255.35 | 3.098 |
| 5V93_B<br>115 | **10300**<br>**50000** | **1312.98**<br>**5900.58** | **3.118**<br>**3.771** | 7656 | 1394.24 | 3.144 |
| 5XBL_B | **3298** | **238.168** | **2.376** | 719 | 128.226 | 2.108 |

| | | | | | | |
|---|---|---|---|---|---|---|
| 98 | **2816** | **83.3305** | **1.921** | | | |
| 6AEB_E 100 | **2643** **2347** | **194.632** **70.6978** | **2.289** **1.849** | 713 | 147.608 | 2.169 |
| 6AGB_A 369 | **286** **265** | **235.667** **106.009** | **2.372** **2.025** | 230 | 173.75 | 2.239 |
| 6MJ0_B 104 | **720** **144** | **114.497** **128.702** | **2.058** **2.109** | 284 | 107.904 | 2.033 |
| 6AHU_A 341 | **2868** **1303** | **1493.51** **870.349** | **3.174** **2.939** | 2748 | 1157.62 | 3.06 |
| 6FT6_2 162 | **3255** **50000** | **484.737** **1483.01** | **2.686** **3.171** | 3810 | 995.342 | 2.997 |
| 6OSI_RB 122 | **10127** **50000** | **1421.84** **5321.56** | **3.153** **3.726** | 6015 | 736.649 | 2.867 |
| 6QX9_1 164 | **12777** **16751** | **6120.5** **6513.45** | **3.787** **3.814** | 14979 | 6620.29 | 3.821 |
| 6WLM_A 171 | **3715** **4181** | **2301.13** **1404.8** | **3.361** **3.147** | 4895 | 1206 | 3.08 |
| 6WLK_A 130 | **41** **101** | **37** **95** | **1.568** **1.978** | 13 | 7 | 0.845 |
| 6ZVK_E1 153 | **200** **235** | **154.253** **185.402** | **2.188** **2.268** | 222 | 160.358 | 2.205 |
| 6UES_A 119 | **4363** **4088** | **1340.81** **1196.01** | **3.127** **3.078** | 2058 | 308.615 | 2.489 |
| 6K4S_A 83 | **759** **1466** | **78.422** **59.382** | **1.894** **1.774** | 371 | 44.4162 | 1.647 |
| 6S0X_B 115 | **11196** **50000** | **2536.79** **4841.23** | **3.404** **3.685** | 7562 | 1954.08 | 3.290 |
| 7OZS_3 119 | **31010** **50000** | **2653.83** **12065.6** | **3.424** **4.082** | 31534 | 2107.23 | 3.323 |
| 7OBQ_1 249 | **8955** **50000** | **3550.35** **1829.79** | **3.550** **3.262** | 7973 | 3324.03 | 3.521 |
| 7RYG_B 115 | **14013** **50000** | **3589.05** **4761.42** | **3.554** **3.678** | 8330 | 2172.88 | 3.337 |
| 7SAM_A 171 | **317** **429** | **220.69** **350.249** | **2.344** **2.544** | 851 | 59.714 | 1.776 |
| 7S4V_B 98 | **3306** **2850** | **256.462** **94.3385** | **2.409** **1.974** | 708 | 150.375 | 2.177 |

| | | | | | | |
|---|---|---|---|---|---|---|
| 7URM_V<br>90 | **16825**<br>**50000** | **3550.57**<br>**4439.71** | **3.550**<br>**3.647** | 19050 | 4432.8 | 3.647 |
| 7SHX_A<br>94 | **31**<br>**345** | **27**<br>**312.375** | **1.431**<br>**2.495** | 26 | 19.78 | 1.296 |
| 7S38_R<br>102 | **2151**<br>**1654** | **269.382**<br>**77.856** | **2.430**<br>**1.891** | 672 | 167.486 | 2.224 |
| 7LMA_B<br>159 | **34**<br>**59** | **21.977**<br>**39.815** | **1.342**<br>**1.600** | 21 | 15.15 | 1.180 |
| 7UVT_A<br>387 | **5594**<br>**8769** | **4089.73**<br>**5087.44** | **3.612**<br>**3.706** | 4739 | 2704.08 | 3.432 |
| 7X34_3<br>130 | **9521**<br>**50000** | **2268.46**<br>**4561.29** | **3.356**<br>**3.659** | 9092 | 2131.55 | 3.328 |
| 7WB1_D<br>121 | **85**<br>**124** | **17.1103**<br>**39.5139** | **1.233**<br>**1.597** | 38 | 15 | 1.17 |
| 7VW3_B<br>104 | **192**<br>**377** | **46.5801**<br>**215.621** | **1.668**<br>**2.334** | 82 | 34.9187 | 1.543 |
| 7YR7_A<br>118 | **1332**<br>**759** | **747.781**<br>**585.189** | **2.873**<br>**2.767** | 1290 | 616.349 | 2.789 |
| 7YOJ_B<br>174 | **133**<br>**680** | **87.4373**<br>**541.583** | **1.942**<br>**2.734** | 31 | 22.4167 | 1.351 |
| 7JRS_B<br>129 | **64**<br>**582** | **52**<br>**549.933** | **1.716**<br>**2.740** | 40 | 27.15 | 1.433 |
| 7ELE_G<br>88 | **4068**<br>**2013** | **1431.93**<br>**637.315** | **3.156**<br>**2.804** | 2494 | 1441.02 | 3.158 |
| 8G1I_B<br>98 | **3408**<br>**2945** | **278.372**<br>**113.585** | **2.445**<br>**2.055** | 718 | 187.579 | 2.273 |
| 8I0P_F<br>107 | **15928**<br>**50000** | **123.355**<br>**13876.6** | **2.091**<br>**4.142** | 6755 | 5.3549 | 0.728 |
| 8SA6_A<br>210 | **6800**<br>**50000** | **5365.14**<br>**27136.5** | **3.729**<br>**4.434** | 6148 | 4822.42 | 3.683 |
| 8BD5_B<br>256 | **249**<br>**501** | **93.061**<br>**257.181** | **1.969**<br>**2.410** | 156 | 84.888 | 1.928 |
| 8QO2_A<br>222 | **44**<br>**73** | **30.667**<br>**60.75** | **1.487**<br>**1.784** | 218 | 41.814 | 1.621 |
| 8VCI_A<br>118 | **13**<br>**62** | **6**<br>**36.6227** | **0.778**<br>**1.564** | 56 | 33.75 | 1.528 |
| 8KAG_A | **3437** | **353.177** | **2.547** | 878 | 284.734 | 2.454 |

| | | | | | | |
|---|---|---|---|---|---|---|
| 98 | **2816** | **177.52** | **2.249** | | | |
| 9ENF_C<br>94 | **26299**<br>**50000** | **2844.66**<br>**2830.63** | **3.454**<br>**3.451** | 11034 | 1511.48 | 3.179 |
| 9G7C_A<br>224 | **3010**<br>**5621** | **978.731**<br>**999.313** | **2.990**<br>**2.999** | 793 | 169.275 | 2.228 |
| 7JRT_B<br>134 | **1**<br>**6574** | **1**<br>**1735.96** | **0**<br>**3.239** | 0 | 0 | 0 |
| 8BTZ_A<br>238 | **29**<br>**263** | **28**<br>**195.242** | **1.447**<br>**2.291** | 89 | 53.4857 | 1.728 |
| 8T2O_R<br>90 | **13**<br>**157** | **9**<br>**153** | **0.954**<br>**2.184** | 8 | 6 | 0.778 |
| 8BU8_A<br>354 | **110**<br>**666** | **92.1006**<br>**444.076** | **1.964**<br>**2.647** | 43 | 42 | 1.623 |
| 8T2P_A<br>135 | **83**<br>**516** | **63.1167**<br>**484.186** | **1.800**<br>**2.685** | 38 | 21.6786 | 1.336 |
| **Mean $\log_{10}$(Neff)** | | | **2.567**<br>**2.617** | | | 2.378 |
| **Median $\log_{10}$(Neff)** | | | **2.879**<br>**2.783** | | | 2.364 |

**Table S5. Comparison of the depths of the MSAs generated for the benchmarking dataset SET-2.**

| | rMSA (with elDORS)<br>**RNAcmap3 (with elDORS)** | | | Standard rMSA (RNAcentral+nt) | | |
|---|---|---|---|---|---|---|
| Sequence_ID<br>_Chain_ID<br>Length | Number of sequences | Neff | $\log_{10}$(Neff) | Number of homologs | Neff | log(Neff) |
| CASP15 | | | | | | |
| 7QR4_B<br>69 | **1245**<br>**546** | **348.008**<br>**435.441** | **2.542**<br>**2.639** | 3780 | 213.76 | 2.329 |
| 8FZA_A<br>30 | **344**<br>**612** | **64.582**<br>**263.267** | **1.810**<br>**2.420** | 166 | 10.49 | 1.021 |
| CASP16 | | | | | | |
| 9CFN_A<br>59 | **13**<br>**177** | **7.141**<br>**163.105** | **0.870**<br>**2.212** | 17 | 3.104 | 0.492 |
| 9C2K_3<br>72 | **2212**<br>**3660** | **62.294**<br>**1124.94** | **1.794**<br>**3.051** | 1902 | 30.647 | 1.486 |

| | | | | | | |
|---|---|---|---|---|---|---|
| 9DCF_2<br>90 | **180**<br>**390** | **44.738**<br>**216.734** | **1.651**<br>**2.336** | 2972 | 94.617 | 1.976 |
| 9ISV_A<br>580 | **1435**<br>**1330** | **772.813**<br>**350.649** | **2.888**<br>**2.544** | 647 | 252.997 | 2.403 |
| 9BZC_A<br>89 | **3519**<br>**2293** | **742.074**<br>**290.577** | **2.870**<br>**2.463** | 2102 | 546.786 | 2.738 |
| 9ELY_A<br>205 | **2950**<br>**5481** | **920.272**<br>**871.172** | **2.964**<br>**2.940** | 1185 | 198.567 | 2.297 |
| 9B0L_5<br>247 | **56**<br>**119** | **41**<br>**84.6608** | **1.613**<br>**1.927** | 246 | 141.152 | 2.149 |
| R0252<br>520 | **58**<br>**108** | **33**<br>**59.131** | **1.518**<br>**1.772** | 107 | 22.3096 | 1.348 |
| R0254<br>413 | **4127**<br>**5954** | **2579.65**<br>**2488.79** | **3.412**<br>**3.396** | 278 | 94.3806 | 1.975 |
| R1241<br>480 | **6011**<br>**50000** | **3822.75**<br>**276.273** | **3.582**<br>**2.441** | 4299 | 2306.89 | 3.363 |
| R1255<br>124 | **18**<br>**302** | **11**<br>**271.23** | **1.041**<br>**2.433** | 222 | 9.4106 | 0.974 |
| R1260<br>387 | **5594**<br>**8769** | **4089.73**<br>**5087.44** | **3.612**<br>**3.706** | 4685 | 2887.25 | 3.460 |
| R1271<br>77 | **21847**<br>**50000** | **1976.73**<br>**4214.97** | **2.296**<br>**3.625** | 17252 | 1416.65 | 3.151 |
| R1286<br>526 | **162**<br>**147** | **117.819**<br>**87.2647** | **2.071**<br>**1.941** | 71 | 34.3667 | 1.536 |
| R1289<br>284 | **4332**<br>**31824** | **2953.04**<br>**4296.92** | **3.740**<br>**3.633** | 4540 | 2277.86 | 3.357 |
| R1293<br>82 | **2**<br>**60** | **2**<br>**60** | **0.301**<br>**1.778** | 25 | 7.6697 | 0.885 |
| **Mean $log_{10}$(Neff)** | | | **2.199**<br>**2.652** | | | 2.052 |
| **Median $log_{10}$(Neff)** | | | **2.183**<br>**2.452** | | | 2.062 |

**Table S7.** Sequences, tertiary structure-derived (ground truth) secondary structures, and the predicted secondary structures (with RNAfold and MXfold2) for the benchmarking dataset SET-3. The ground truth secondary structures are plotted using VARNA.

| Sequence_ID_ | Sequence | 2D |
|---|---|---|

| Chain_ID<br>Length | | |
|---|---|---|
| 2DU4_C<br>71 | GCCAGGGUGGCAGAGGGGCUUUGCG<br>GCGGACUGCAGAUCCGCUUUACCCC<br>GGUUCGAAUCCGGGCCCUGGC | 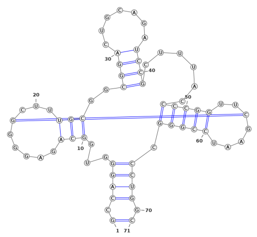<br>(.((((...(((.....[...)))..((((.......))))......(((((..]....))))).)))).)<br><br>**RNAfold**<br>(((((((..(((.........))).(((((.......))))).....(((((.......))))))))))))<br>**MXfold2**<br>((((((((((.((((.((.(((((((....))))))).)).)))).))((((.......)))))))))))) |
| 2QUS_A<br>69 | GGGAGCCCUGUCACCGGAUGUGCUU<br>UCCGGUCUGAUGAGUCCGUGAGGAC<br>AAAACAGGGCUCCCGAAUU | 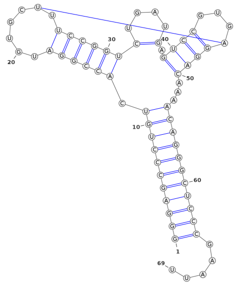<br>((((((((((((.((((((.....[.))))))(....).((((...]))))...))))))))))))).....<br>**RNAfold**<br>((((((((((((.((((((.......))))))).......((((....))))...))))))))))))).....<br><br>**MXfold2**<br>((((((((((((.((((((.......))))))).......((((....))))...))))))))))))))..... |
| 4N0T_B<br>72 | GGUCAAUUUGAAACAAUACAGAGAU<br>GAUCAGCGGUUCCCCUGCAUAAGGA<br>UGAACCGUUUUACAAAGAGACC | 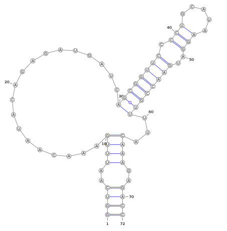<br>.(((..((((..................((((((((.((.-------))..))))))))...))))..)))-<br>**Approximated**<br>((((..((((..................((((((((.(((.....)))..))))))))...))))..))))<br><br>**RNAfold**<br>(((((.((((........))))..)))))((((((((.(((.....)))..))))))))............<br><br>**MXfold2**<br>((((..((((........((....))...((((((((.(((.....)))..))))))))...))))..)))) |

| | | |
|---|---|---|
| 5LYU_A<br>58 | GGGGAUCUGUCACCCCAUUGAUCGCCUUCGGGCUGAUCUGGCUGGCUAGGCGGGUCCC | 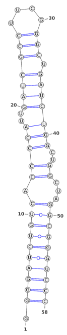<br>**.((((((((((.(((((..(((((((....))).)))))))..))...))))))))))**<br><br>**RNAfold**<br>**.((((((((((.(((((..(((((((....))).)))))))..))...))))))))))**<br>**MXfold2**<br>**.((((((((((.(((((..(((((((....))).)))))))..))...))))))))))** |
| 6OL3_C<br>112 | GGACCUCGCAAGGGUAUCAUGGCGGACGACCGGGGUUCGAACCCCGGAUCCGGCCGUCCGCCGUGAUCCAUGCGGUUACCGCCCGCGUGUCGAACCCAGGUGUGCGAGGUCC | 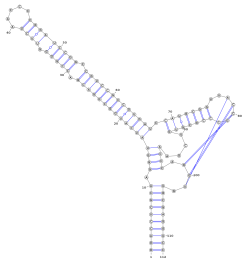<br>**((((((((((.(((((((((((((((((.(((((-((((.....))))))))).))))))))))))).((((((**<br>**(..[[[..)))))))....)))).]]].))))))))))**<br>**Approximated**<br>**((((((((((.(((((((((((((((((.(((((.((((.....))))))))).))))))))))))).((((((**<br>**(..[[[..)))))))....)))).]]].))))))))))**<br><br>**RNAfold**<br>**((((((((((.(((((((((((((((((.(((((.((((.....))))))))).))))))))))))).((((((**<br>**(.......)))))))....)))).....))))))))))**<br>**MXfold2**<br>**((((((((((.(((((((((((((((((.(((((((....))))))).......))))))))))))).(((((((**<br>**.......)))))))....)))).....))))))))))** |
| 1XJR_A<br>47 | GGAGUUCACCGAGGCCACGCGGAGUACGAUCGAGGGUACAGUGAAUU | 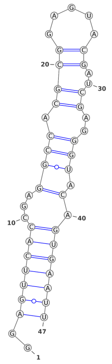<br>**..(((((((...((((.((((.....))..))..))).).)))))))**<br><br>**RNAfold** |

| | | |
|---|---|---|
| | | ..(((((((.(..(((.((((.....))..))..))).).)))))))<br>**MXfold2**<br>...((((((....(((.((((.....))..))..)))...)))))). |
| 6MWN_A<br>92 | GGCAAACAUCAUUUGGCCUUAAAUG<br>GGAUUCUGUGAGAGGGGAUCCCUCC<br>AUUGACAGCUGGACUGUUCUUUGGG<br>GCCUUAUGUGGUGUUUG | 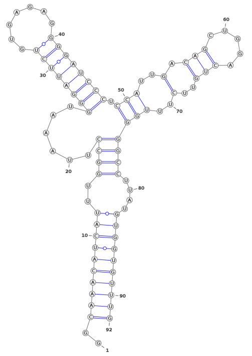<br>..((((((((((..((((......((((((((.......)))))))).(((..(.(((.....)))..)..))))))).<br>...))))))))))<br><br>**RNAfold**<br>..((((((((((..(((((((((.((((((((.......))))))))......(((((.....))))).))))))))<br>)....))))))))))<br>**MXfold2**<br>..((((((((((..(((((((((.((((((((.......))))))))......(((((.....))))).))))))))<br>)....)))))))))) |
| 6MJ0_B<br>104 | UAAGUUCUCGAUCUUUAAAAUC<br>GUUAGCUCGCCAGUUAGCGAGG<br>UCUGCGAAAGCAGAUAAUCGGG<br>UGCAACUCCCGCCCUUUCUCCGA<br>GGGUCAUCGGAACCA | 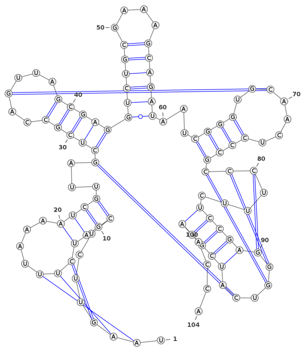<br>-[[[....((((]]]....))))...[(((((..[...)))))((((((....))))))...((((..]....))))[[<br>[....((((((]]].].))))))..--<br>**Approximated**<br>.[[[....((((]]]....))))...[(((((..[...)))))((((((....))))))...((((..]....))))[[<br>[....((((((]]].].)))))).....<br>**RNAfold**<br>...((((.((((........(((((((((....))))))))).((((((....))))))....(((.......)))(((<br>(((.....)))))).)))))))).<br>**MXfold2**<br>...((((.((((........(((((((((....))))))))).((((((....))))))....(((.......)))(((<br>(((.....)))))).))))))).. |

| | | |
|---|---|---|
| 5T83_A<br>95 | GUCUAAAGUUUGCUAGGGUUCC<br>GCGUCAUAGGUGGUCUGGUCCA<br>AGAGCAAACGGCUUUCACAAAG<br>CCACACGGAAGGAUAAAAGCCU<br>GGGAGAU | 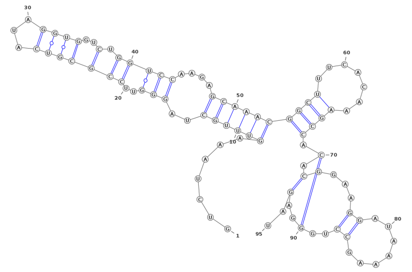<br>**-----..((((((..(((..(((((((...))))..).)))))....))))))((((.......)))).[.(....((.**<br>**......))..]..)..**<br>**Approximated**<br>**.......((((((..(((..(((((((...))))..).)))))....))))))((((.......)))).[.(....((...**<br>**....))..]..)..**<br>**RNAfold**<br>**.......((((.(((((.((((...((....(((((...((...(((((.....))))).))...)))))...)).)))..**<br>**..).))))).))))**<br>**MXfold2**<br>**((((...(((((((.(((..((((.......))))).....)))...)))))))((((((...))))))).......(((..**<br>**.....)))...))))** |
| 4V2S_Q<br>65 | UUCCGAUGUAGACCCGUCcUCCUUC<br>GCCUGCGUCACGGGUCCUGGUUAGA<br>CGCAGGCGUUUUCUG | 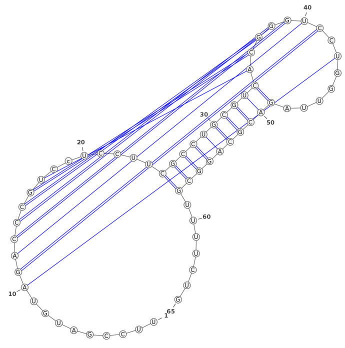<br>**-----....[[[[[[[[..--...((((((((((]]]]]]].].....)))))))))).......**<br>**Approximated**<br>**.........[[[[[[[[..--...((((((((((]]]]]]].].....)))))))))).......**<br><br>**RNAfold**<br>**....((((......))))......(((((((((((........))..))))))))).......**<br>**MXfold2**<br>**......................((((((((((((.(....).))..)))))))))).......** |
| 6JDV_B<br>147 | GGUCACUCUGCUAUUUAACUUUACG<br>UUGUAGCUCCCUUUCUCAUUUCGGA<br>AACGAAAUGAGAACCGUUGCUACAA<br>UAAGGCCGUCUGAAAAGAUGUGCCG<br>CAACGCUCUGCCCCUUAAAGCUCCU<br>GCUUUAAGGGGCAUCGUUUAUC | 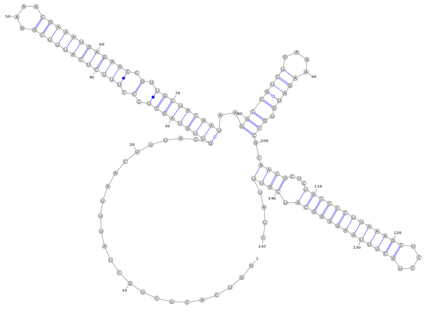<br>**......................(((((((((.(.((((((((((((....))))))))))).).)))))))))..((**<br>**(((((((....))))).)))..((((...(((((((((((((....)))))))))))).))))....** |

<table>
<tr><td></td><td></td><td>RNAfold<br>........(((.............((((((((..(..((((((((((((....))))))))))))..)..))))))))..((<br>((((((....))))).))))))(((...(((((((((((((....))))))))))))).))).....<br>MXfold2<br>......................((((((((..(..((((((((((((....))))))))))))..)..))))))))..(((<br>(((((....))))).)))...(((...(((((((((((((....))))))))))))).))).....</td></tr>
<tr><td>4PQV_A<br>68</td><td>GGGUCAGAUCGGCGAAAGUCGCCAC<br>UUCGCCGAGGAGUGCAAUCUGUGAG<br>GCCCCAGGAGGACUGGGU</td><td>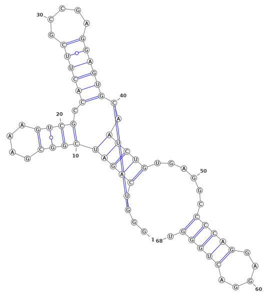<br>..[.(((((((((....)))).((((((.....))))))].)))))......(((((.....))))).<br>RNAfold<br>..........(((((...)))))......(((..(.((......)).)..)))(((((.....))))).<br>MXfold2<br>........((((((.((((....))))))))))..................(((((.....))))).</td></tr>
<tr><td>4FRN_A<br>102</td><td>GGGCUAAAAGCAUGGUGGGAAAGUG<br>ACGUGUAAUUCGUCCACAUUACUUG<br>AUACGGUUAUACUCCGAAUGCCACC<br>UAGCCCAAAGUAGAGCAAGGAGACU<br>CA</td><td>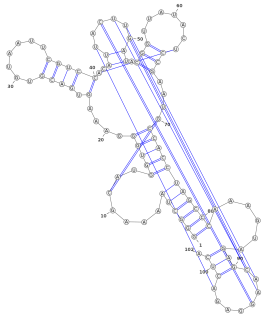<br>((((((....{..(((((....(.((((.......))))){.(([.[[[[)).(((......}))))...}))))))))<br>))).......(((]]]]...]))).<br><br>RNAfold<br>((((((.......(((((.(..(.((.(((((((.(((...((.....))..)))))))))))))...).))))))))<br>))).......(((........))).<br>MXfold2<br>((((((.......(((((.(((((((.(((.........))).))))))....(((.......)))..).))))))))))<br>).......................</td></tr>
<tr><td>6QN3_A<br>50</td><td>CGUUCACCCUUCGGGGCGCAUGAAA<br>UGGGAGUAGGGAACGGGAUUCUCAU</td><td>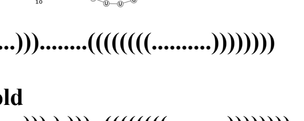<br>......(((....)))........((((((((..........))))))))<br><br>RNAfold<br>(((.(.((((....)))).).)))..((((((((..........))))))))<br>MXfold2<br>(((.(.((((....)))).).)))...........................</td></tr>
</table>

| | | |
|---|---|---|
| 4L81_A<br>97 | GGAUCACGAGGGGGAGACCCCGGCA<br>ACCUGGGACGGACACCCAAGGUGCU<br>CACACCGGAGACGGUGGAUCCGGCC<br>CGAGAGGGCAACGAAGUCCGUA | 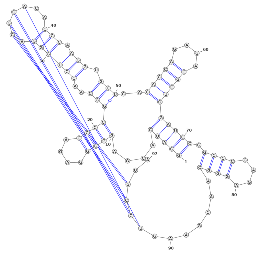<br>**(((((....((((....))))(((..((((((.[[[[..))).))).)))..(((((....)))))))))).((((.<br>...)))).......]]]]..**<br><br>**RNAfold**<br>**(((((....((((....))))((((.(((((((.......))).)))))))..(((((....)))))))))).((((..<br>..)))).(((.....))).**<br>**MXfold2**<br>**(((((....((((....))))(((.((((((((.......)))).))))))..(((((....)))))))))).((((..<br>..)))).............** |
| 4ENA_A<br>52 | GGGCGAUGAGGCCCGCCCAAACUGC<br>CCUGAAAAGGGCUGAUGGCCUCUAC<br>UG | 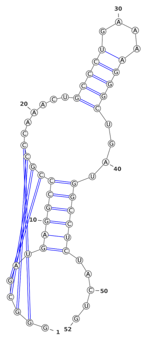<br>**.[[[[..((((((]]]]......(((((....)))))....))))))).....**<br><br>**RNAfold**<br>**((((......))))(((......(((((....)))))....)))........**<br>**MXfold2**<br>**((((......))))(((......(((((....)))))....)))........** |
| 4XWF_A<br>64 | GGGUCGUGACUGGCGAACAGGUGGG<br>AAACCACCGGGGAGCGACCCUUGCC<br>GCCCGCCUGGGCAA | 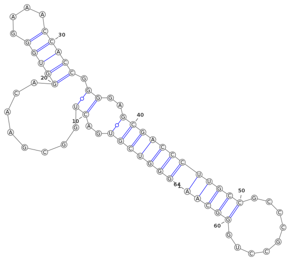<br>**(((((((..((........(((((....))))).))..)))))))(((((.........)))))**<br><br>**RNAfold**<br>**(((((((..((........(((((....))))).))..))))))).....((((....))))..**<br>**MXfold2**<br>**(((((((..((........(((((....))))).))..))))))).....((((....))))..** |

| | | |
|---|---|---|
| 5O69_A<br>41 | CCGGACGAGGUGCGCCGUACCCGGU<br>CAGGACAAGACGGCGC | 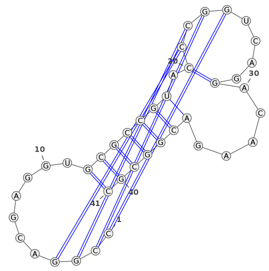<br>**[[[[.......(((((((.(]]]]....).....)))))))**<br><br>**RNAfold**<br>**((......)).(((((((..((.....)).....)))))))**<br>**MXfold2**<br>**...........(((((((..((.....)).....)))))))** |
| 3Q3Z_A<br>75 | GCGCGGAAACAAUGAUGAAUGGGUU<br>UAAAUUGGGCACUUGACUCAUUUUG<br>AGUUAGUAGUGCAACCGACCGUGCU | 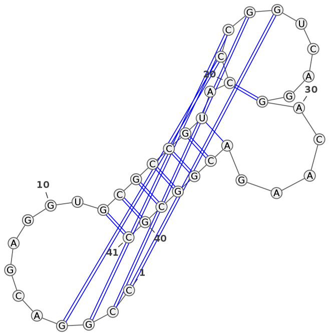<br>**((((((..((......(((((((((.....{[[[[[[[.)))))))))...))....]]]]]}.]]..)))))).**<br><br>**RNAfold**<br>**((((((((((.((.....))..))))...((((((((((((((((...)))))))..)))))..)))))))))).**<br>**MXfold2**<br>**((((((...((((................)))).(((((((((((((...)))))))..)))))......)))))).** |
| 6FZ0_A<br>54 | AGGCGCAUUUGAACUGUAUUGUACG<br>CCUUGCAUAAAGCAAAAGUACUAAA<br>AAAA | 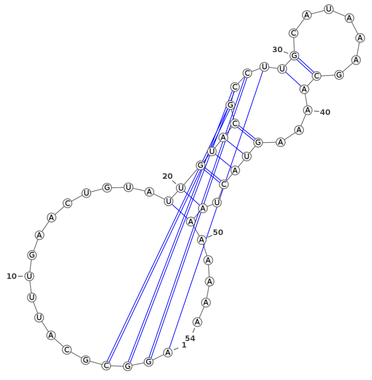<br>**[[[[..............((((((]]]]((..----.))...)))).))...-.**<br>**Approximated**<br>**[[[[..............((((((]]]]((.......))...)))).)).....**<br><br>**RNAfold**<br>**.((((.((............)).))))((((.....))))..............** |

| | | |
|---|---|---|
| | | **MXfold2**<br>**...................((((...((((.....))))..))))........** |
| 3ADB_C<br>92 | GGCCGCCGCCACCGGGGUGGUCCCC GGGCCGGACUUCAGAUCCGGCGCGC CCCGAGUGGGGCGCGGGGUUCAAUU CCCCGCGGCGGCCGCCA | 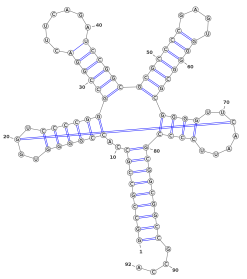<br>**(((((((((..((((((..[.))))))((((((.......))))))(((((((....)))))))((((..]....))**<br>**)))))))))))....**<br>**RNAfold**<br>**(((((((((..((((((....))))))((((((.......))))))(((((((....)))))))((((.......))**<br>**)))))))))))....**<br>**MXfold2**<br>**(((((((((..((((((....))))))((((((.......))))))((((((......))))))((((.......)))**<br>**))))))))))....** |
| 4ATO_G<br>34 | AAAUUGGUGUAACCUUACCGUAGUA GGUGCUAAA | 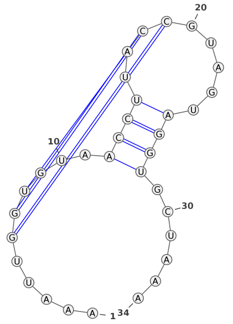<br>**.....[[[...((((.]]].....)))).......**<br><br>**RNAfold**<br>**...(((((...((((.((....))))))))))))).**<br>**MXfold2**<br>**...(((((...(((.(((....)))))))))))).** |
| 6CU1_A<br>81 | GGGUGGCUCUACAUUUGUUGCGGGU UCGAGACCCGUCAGAGCGAAAGCUC UGUAGCUCAAUGGUAGAGCGGUGUA GUCACC | 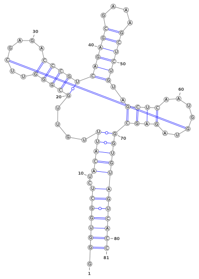<br>**.(((((((.-(((((....(((((..[....)))))((((((....))))))..((((....]..))))))))))))**<br>**))))**<br>**Approximated**<br>**.(((((((..(((((....(((((..[....)))))((((((....))))))..((((....]..))))))))))))))))** |

| | | |
|---|---|---|
| | | )))<br>**RNAfold**<br>**.....(((((((..(((..(((((((...)))))))((((((....)))))).....)))..)))))))((((....)**<br>**)))**<br>**MXfold2**<br>**.(((((..((((((((....((((((.......)))))((((((....))))))..((((.......))))))))))))**<br>**)))** |
| 6JQ5_A<br>82 | UUACUGUGAGAAUCAGUAACAAACA UGUGGGGCUUAUAUCUAAUCGAAAG AUUAGUAUUAGUGCAGACGUUAAAA CCAUGUC | 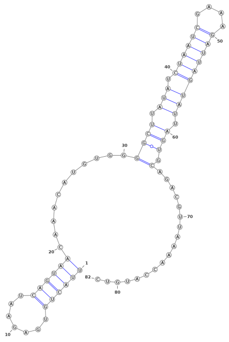<br>**((((((.......))))))...........((((.(((.((((((....))))))))).))).)................**<br><br>**RNAfold**<br>**((((((.......))))))......((((.(.(((.(((.((((((....))))))))).))).)..))))..........**<br>**..**<br>**MXfold2**<br>**.((((((.......)))))....(((((..(.(((.(((.((((((....))))))))).))).)...........))))**<br>**).** |
| 6UFJ_A<br>51 | ACUCGUUUGAGCGAGUAUAAACAGC UGGUUAAGCUCAAAGCGGAGAGCAG A | 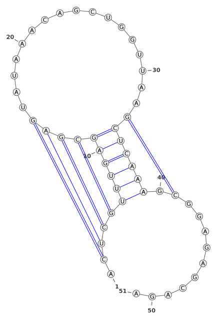<br>**.[[[[((((((([]]]].................[))))))).]..........-**<br>**Approximated**<br>**.[[[[((((((([]]]].................[))))))).]..........**<br>**RNAfold**<br>**((((((....))))))........(((.((..(((...)))...)).))).**<br>**MXfold2**<br>**.((((((....))))))..................((((........))))...** |

| | | |
|---|---|---|
| 4RZD_A<br>101 | GAGCAACUUAGGAUUUUAGGCUCCCCGGCGUGUCUCGAACCAUGCCGGGCCAAACCCAUAGGGCUGGCGGUCCCUGUGCGGUCAAAAUUCAUCCGCCGGAG | 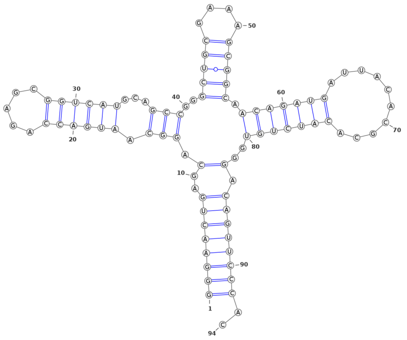<br>**[[[[......((((((...]]]](((((((((.--.....))))))))).....((((((((((........)))))))<br>.)))..))))))...........**<br>**Approximated**<br>**[[[[......((((((...]]]](((((((((........)))))))))....((((((((((........))))))).<br>)))..))))))...........**<br><br>**RNAfold**<br>**((((...............)))).((((((((........))))))))((((..((....))..)))).....(((.((((<br>...........)))))))..**<br>**MXfold2**<br>**((((...............))))(((((((((........)))))))))).....(((...)))(((((((...........<br>..........)))))))..** |
| 4RUM_A<br>94 | GGGAACUGAGCAGGCAAUGACCAGAGCGGUCAUGCAGCCGGGCUGCGAAAGCGGCAACAGAUGAUUACACGCACAUCUGUGGGACAGUUCCCAC | 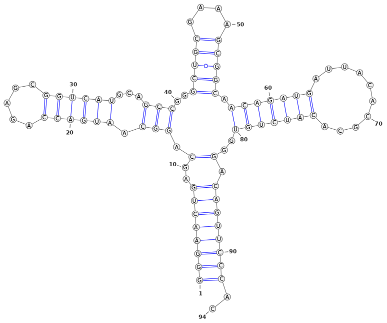<br>**((((((((..(.(((.((((((.....))))))...)))..(((((....))))).(((((((..-.......)))))<br>))..).))))))))..**<br>**Approximated**<br>**((((((((..(.(((.((((((.....))))))...)))..(((((....))))).(((((((..........))))))<br>)..).))))))))..**<br><br>**RNAfold**<br>**((((((((..(.(((.((((((.....))))))...)))..(((((....))))).(((((((..........))))))<br>).)..))))))))..**<br>**MXfold2**<br>**((((((((....(((.((((((.....))))))...)))..(((((....))))).(((((((..........))))))<br>)....))))))))..** |

| | | |
|---|---|---|
| 3LWO_D<br>58 | GGGCCACGGAAACCGCGCGCGGUGA<br>UCAAUGAGCCGCGUUCGCUCCCGUG<br>GCCCACAA | 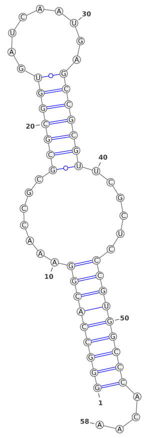<br>(((((((((.......(((((((.........)))))))......)))))))))....<br><br>**RNAfold**<br>(((((((((....((.(((((((.........))))))).))...)))))))))....<br>**MXfold2**<br>(((((((((..(.((.(((((((.........))))))).)).).))))))))).... |
| 6TFE_A<br>52 | GGCUUCAACAACCCCGUAGGUUGGG<br>CCGAAAGGCAGCGAAUCUACUGGAG<br>CC | 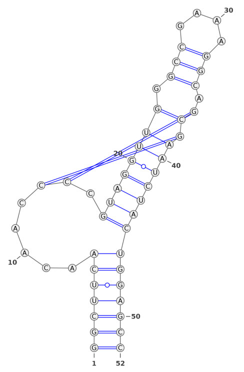<br>(((((((.....[[.((((((((.(((....))).])]))))))))))))))<br><br>**RNAfold**<br>(((((((........(((((((.((((....)))..).))))))))))))))<br>**MXfold2**<br>(((((((.(((((.....))))).(((....)))...........))))))) |

**Table S7. Comparison of the depths of the MSAs generated for the benchmarking dataset SET-3.**

**A.** Comparison based on the number of effective sequences in the MSA and the F1-score for RNA secondary structure prediction

| PDB_ID<br>_Chain ID | Description | **RNAcmap3-elDORS**<br>**(RNAcmap3-elDORS-PK)**<br>**RNAcmap3-mx-elDORS** | RNAcmap3-MARS<br>**RNAcmap2-MARS**<br>*Rfam* |
|---|---|---|---|

| | | (RNAcmap3-mx-elDORS-PK) | | | | | | |
|---|---|---|---|---|---|---|---|---|
| Rfam-mapped sequences | | No. of sequences | Neff | $\log_{10}$ (Neff) | F1-score | [a]Neff | $\log_{10}$(Neff) | [a]F1-score |
| **2DU4_C** | **RF00005 tRNA** | **12561**<br>**50000** | **868.316**<br>**5310.43** | **2.938**<br>**3.725** | **0.683**<br>**(0.700)**<br>**0.878**<br>**(0.850)** | 712.4<br>**59.0**<br>*3037.1* | 2.853<br>**1.771**<br>*3.482* | 0.556<br>**0.704**<br>*0.778* |
| **2QUS_A** | **RF00008 Hammerhead ribozyme** | **2399**<br>**2399** | **1223.27**<br>**1223.27** | **3.087**<br>**3.087** | **0.826**<br>**0.844**<br>**0.826**<br>**0.844** | 1340.9<br>**149.2**<br>*171.0* | 3.127<br>**2.174**<br>*2.233* | 0.792<br>**0.792**<br>*0.604* |
| **4N0T_B** | **RF00026 U6 spliceosomal** | **2710**<br>**4987** | **206.341**<br>**291.68** | **2.269**<br>**2.464** | **0.465**<br>**0.884** | 158.2<br>**227.9**<br>*2412.8* | 2.199<br>**2.358**<br>*3.382* | 0.465<br>**0.439**<br>*0.000* |
| **5LYU_A** | **RF00100 7SK** | **12326**<br>**12326** | **179.385**<br>**179.385** | **2.254**<br>**2.254** | **0.927**<br>**0.927** | 192.9<br>**45.9**<br>*1832.6* | 2.285<br>**1.662**<br>*3.263* | 0.844<br>**0.622**<br>*0.844* |
| **6OL3_C** | **RF00102 VA** | **656**<br>**387** | **554.379**<br>**293.802** | **2.744**<br>**2.468** | **0.881**<br>**(0.914)**<br>**0.667**<br>**(0.691)** | 404.3<br>**65.4**<br>*29.7* | 2.607<br>**1.816**<br>*1.472* | 0.782<br>**0.782**<br>*0.409* |
| **1XJR_A** | **RF00164 Coronavirus 3′ stem-loop-II-like motif** | **502**<br>**653** | **273.672**<br>**350.133** | **2.437**<br>**2.544** | **0.933**<br>**0.867** | 204.1<br>**21.7**<br>*4.0* | 2.309<br>**1.336**<br>0.602 | 0.824<br>**0.588**<br>*0.412* |
| **6MWN_A** | **RF00228 Hepatitis A virus IRES** | **144**<br>**144** | **114.644**<br>**114.644** | **2.06**<br>**2.06** | **0.814**<br>**0.814** | 249.0<br>**633.2**<br>*1* | 2.396<br>**2.801**<br>*0* | 0.716<br>**0.716**<br>*0.090* |
| **6MJ0_B** | **RF00390 UPSK** | **144**<br>**144** | **128.702**<br>**128.702** | **2.109**<br>**2.109** | **0.364**<br>**(0.310)**<br>**0.364**<br>**(0.310)** | 135.8<br>**26.8**<br>***1*** | 2.133<br>**1.428**<br>*0* | 0.316<br>**0.306**<br>*0.066* |
| **5T83_A** | **RF00442 Guanidine-I riboswitch** | **441**<br>**3186** | **137.036**<br>**357.037** | **2.137**<br>**2.552** | **0.182**<br>**(0.185)**<br>***0.545***<br>***(0.556)*** | 169.4<br>**300.1**<br>*152.7* | 2.229<br>**2.477**<br>*2.184* | 0.247<br>**0.250**<br>*0.556* |
| **4V2S_Q** | **RF00505 RydC** | **1792**<br>**787** | **1391.83**<br>**591.301** | **3.143**<br>**2.772** | **0.513**<br>**(0.645)**<br>**0.513**<br>**(0.645)** | 1271.1<br>**65.9**<br>*4* | 3.104<br>**1.819**<br>*0.602* | 0.571<br>**0.526**<br>*0.270* |
| **6JDV_B** | **RF01344** | **703**<br>**814** | **508.903**<br>**530.187** | **2.707**<br>**2.724** | **0.417**<br>**0.667** | 432.3<br>**132.5** | 2.636<br>**2.122** | 0.900<br>**0.880** |

| | | | | | | | | |
|---|---|---|---|---|---|---|---|---|
| | **CRISPR RNA direct repeat element** | | | | | *6.8* | *0.832* | *0.000* |
| **4PQV_A** | **RF01415 Flavivirus 3′ UTR stem loop IV** | **370**<br>**2289** | **318.437**<br>**1668.19** | **2.503**<br>**3.222** | **0.233**<br>**0.238**<br>**0.233**<br>**0.238** | 206.9<br>**9.9**<br>*4.9* | 2.316<br>**0.995**<br>*0.690* | 0.222<br>**0.133**<br>*0.267* |
| **4FRN_A** | **RF01689 AdoCbl variant** | **154**<br>**520** | **93.1793**<br>**116.064** | **1.969**<br>**2.065** | **0.562**<br>**(0.621)**<br>**0.531**<br>**(0.586)** | 107.1<br>**12.3**<br>*60.5* | 2.029<br>**1.089**<br>*1.782* | 0.507<br>**0.427**<br>*0.613* |
| **6QN3_A** | **RF01704 Downstream peptide RNA** | **593**<br>**5064** | **154.442**<br>**3199.69** | **2.189**<br>**3.505** | **0.815**<br>**0.074** | 156.0<br>**38.7**<br>*52.3* | 2.193<br>**1.588**<br>*1.718* | 0.733<br>**0.733**<br>*0.733* |
| **4L81_A** | **RF01725 SAM-I/IV variant riboswitch** | **1330**<br>**1849** | **572.109**<br>**713.376** | **2.757**<br>**2.853** | **0.762**<br>**(0.814)**<br>**0.794**<br>**(0.847)** | 574.0<br>**118.2**<br>*169.8* | 2.759<br>**2.073**<br>*2.229* | 0.750<br>**0.611**<br>*0.500* |
| **4ENA_A** | **RF01734 Fluoride riboswitch** | **3626**<br>**3968** | **362.015**<br>**336.135** | **2.558**<br>**2.526** | **0.437**<br>**(0.500)**<br>**0.437**<br>**(0.500)** | 106.0<br>**52.4**<br>*242.2* | 2.025<br>**1.719**<br>*2.384* | 0.514<br>**0.629**<br>*0.800* |
| **4XWF_A** | **RF01750 ZMP/ZTP riboswitch** | **2379**<br>**2379** | **486.078**<br>**486.078** | **2.687**<br>**2.687** | **0.700**<br>**0.700** | 490.1<br>**262.8**<br>*102.4* | 2.690<br>**2.419**<br>*2.010* | 0.622<br>**0.444**<br>*0.622* |
| **5O69_A** | **RF01763 Guanidine-III riboswitch** | **1785**<br>**3595** | **114.335**<br>**497.208** | **2.058**<br>**2.696** | **0.560**<br>**0.667**<br>**0.560**<br>**0.667** | 112.8<br>**4.1**<br>*5.1* | 2.052<br>**0.613**<br>*0.708* | 0.359<br>**0.513**<br>*0.513* |
| **3Q3Z_A** | **RF01786 Cyclic di-GMP-II riboswitch** | **1042**<br>**966** | **651.943**<br>**652.202** | **2.814**<br>**2.814** | **0.760**<br>**0619**<br>**0.720**<br>**0.619** | 650.6<br>**332.8**<br>*372.1* | 2.813<br>**2.522**<br>*2.571* | 0.576<br>**0.847**<br>*0.847* |
| **6FZ0_A** | **RF01826 SAM-V riboswitch** | **3334**<br>**1637** | **1533.81**<br>**193.518** | **3.186**<br>**2.287** | **0.333**<br>**0.154**<br>**0.462**<br>**0.400** | 2251.0<br>**24.1**<br>*2.9* | 3.352<br>**1.382**<br>*0.462* | 0.238<br>**0.524**<br>*0.300* |
| **3ADB_C** | **RF01852 Selenocysteine transfer** | **6197**<br>**5896**<br>**Different 2D** | **1221.17**<br>**1200.2** | **3.087**<br>**3.079** | **0.952**<br>**0.968**<br>**0.952**<br>**0.968** | 1002.5<br>**56.9**<br>*298.8* | 3.001<br>**1.755**<br>*2.475* | 0.896<br>**0.896**<br>*0.866* |
| **4ATO_G** | **RF02519 ToxI antitoxin** | **159**<br>**163** | **81.386**<br>**88.764** | **1.9105**<br>**1.948** | **0.444**<br>**0.533** | 156.5<br>**15.3** | 2.194<br>**1.185** | 0.364<br>**0.364** |

| | | | | | | | | |
|---|---|---|---|---|---|---|---|---|
| | | | | | 0.333<br>0.400 | *1.2* | 0.079 | *0.091* |
| 6CU1_A | RF02553<br>γ RNA-like | 473<br>9539 | 276.35<br>3776.35 | 2.441<br>3.578 | 0.400<br>0.407<br>0.800<br>0.778 | 381.7<br>**330.6**<br>*84.2* | 2.582<br>**2.519**<br>*1.925* | 0.387<br>**0.476**<br>*0.698* |
| 6JQ5_A | RF02678 Hatchet ribozyme | 206<br>202 | 69.827<br>59.743 | 1.844<br>1.776 | 0.696<br>0.696 | 84.1<br>**9.0**<br>*3.2* | 1.924<br>**0.954**<br>*0.505* | 0.704<br>**0.444**<br>*0.222* |
| 6UFJ_A | RF02679<br>Pistol ribozyme | 256<br>197 | 101.723<br>70.617 | 2.007<br>1.849 | 0.667<br>0.364<br>0.667<br>0.455 | 849.0<br>**47.6**<br>*43.3* | 2.929<br>**1.678**<br>*1.636* | 0.378<br>**0.541**<br>*0.486* |
| 4RZD_A | RF02680<br>PreQ1-III riboswitch | 514<br>572 | 142.408<br>178.996 | 2.154<br>2.253 | 0.419<br>0.310<br>0.452<br>0.345 | 110.7<br>**11.9**<br>*2.8* | 2.044<br>**1.076**<br>*0.447* | 0.353<br>**0.294**<br>*0.261* |
| 4RUM_A | RF02683<br>NiCo riboswitch | 1189<br>1264 | 587.436<br>706.175 | 2.769<br>2.849 | 0.951<br>0.951 | 625.4<br>**61.7**<br>*118.2* | 2.796<br>**1.790**<br>2.073 | 0.866<br>**0.667**<br>*0.627* |
| 3LWO_D | RF02796 Pab160 | 802<br>867 | 689.892<br>757.927 | 2.839<br>2.879 | 0.914<br>0.914 | 592.8<br>**13.5**<br>*3.3* | 2.773<br>**1.130**<br>*0.518* | 0.821<br>**0.667**<br>*0.462* |
| 6TFE_A | RF03013 nadA | 696<br>889 | 256.313<br>500.53 | 2.409<br>2.699 | 0.895<br>0.944<br>0.526<br>0.556 | 190.3<br>**14.0**<br>*28.8* | 2.279<br>**1.146**<br>*1.459* | 0.842<br>**0.632**<br>*0.737* |
| **Mean $\log_{10}$(Neff) and F1-score** | | | | 2.485<br>2.631 | 0.638<br>0.633<br>0.647<br>0.647 | | [b]2.504<br>**1.703**<br>*1.473* | [b]0.591<br>**0.567**<br>*0.472* |
| **Median $\log_{10}$(Neff) and F1-score** | | | | 2.441<br>2.669 | 0.683<br>0.667<br>0.667<br>0.667 | | 2.396<br>**1.719**<br>*1.472* | 0.576<br>**0.588**<br>*0.500* |

[a]Obtained from Chen et al, 2024
[b]Calculated for the 29 sequences.

**B.** Comparison based on Recall (sensitivity), Precision (PPV) for RNA secondary structure prediction (**RNAcmap3-elDORS** **(RNAcmap3-elDORS-PK),** **RNAcmap3-mx-elDORS** **(RNAcmap3-mx-elDORS-PK)**).

| PDB_ID _Chain ID | **Recall (Sensitivity)** | **Precision (PPV)** | **ID** | **Recall (Sensitivity)** | **Precision (PPV)** |
|---|---|---|---|---|---|
| **2DU4_C** | **0.778**<br>**0.824**<br>**1.000**<br>**1.000** | **0.609**<br>**0.609**<br>**0.783**<br>**0.739** | **4ENA_A** | **0.467**<br>**0.636**<br>**0.467**<br>**0.636** | **0.412**<br>**0.412**<br>**0.412**<br>**0.412** |
| **2QUS_A** | **0.870**<br>**0.864**<br>**0.870**<br>**0.864** | **0.870**<br>**0.826**<br>**0.870**<br>**0.826** | **4XWF_A** | **0.737**<br>**0.737**<br>**0.737**<br>**0.737** | **0.667**<br>**0.667**<br>**0.667**<br>**0.667** |
| **4N0T_B** | **0.417**<br>**0.417**<br>**1.000**<br>**1.000** | **0.526**<br>**0.526**<br>**0.792**<br>**0.792** | **5O69_A** | **0.667**<br>**1.000**<br>**0.667**<br>**1.000** | **0.615**<br>**0.615**<br>**0.615**<br>**0.615** |
| **5LYU_A** | **0.864**<br>**0.864**<br>**0.864**<br>**0.864** | **1.000**<br>**1.000**<br>**1.000**<br>**1.000** | **3Q3Z_A** | **0.760**<br>**0.765**<br>**0.720**<br>**0.765** | **0.760**<br>**0.520**<br>**0.720**<br>**0.520** |
| **6OL3_C** | **0.787**<br>**0.841**<br>**0.596**<br>**0.636** | **1.000**<br>**1.000**<br>**0.757**<br>**0.757** | **6FZ0_A** | **0.417**<br>**0.250**<br>**0.500**<br>**0.750** | **0.278**<br>**0.111**<br>**0.333**<br>**0.333** |
| **1XJR_A** | **0.933**<br>**0.933**<br>**0.867**<br>**0.867** | **0.933**<br>**0.933**<br>**0.867**<br>**0.867** | **3ADB_C** | **0.909**<br>**0.938**<br>**0.909**<br>**0.938** | **1.000**<br>**1.000**<br>**1.000**<br>**1.000** |
| **6MWN_A** | **0.828**<br>**0.828**<br>**0.828**<br>**0.828** | **0.800**<br>**0.800**<br>**0.800**<br>**0.800** | **4ATO_G** | **0.571**<br>**1.000**<br>**0.571**<br>**1.000** | **0.364**<br>**0.364**<br>**0.364**<br>**0.364** |
| **6MJ0_B** | **0.375**<br>**0.375**<br>**0.375**<br>**0.375** | **0.353**<br>**0.265**<br>**0.353**<br>**0.265** | **6CU1_A** | **0.407**<br>**0.393**<br>**0.786**<br>**0.778** | **0.407**<br>**0.407**<br>**0.815**<br>**0.778** |
| **5T83_A** | **0.208**<br>**0.217**<br>**0.625**<br>**0.652** | **0.161**<br>**0.161**<br>**0.484**<br>**0.484** | **6JQ5_A** | **0.842**<br>**0.842**<br>**0.895**<br>**0.895** | **0.593**<br>**0.593**<br>**0.630**<br>**0.630** |
| **4V2S_Q** | **0.556**<br>**1.000**<br>**0.556**<br>**1.000** | **0.476**<br>**0.476**<br>**0.476**<br>**0.476** | **6UFJ_A** | **0.818**<br>**0.667**<br>**0.818**<br>**0.833** | **0.562**<br>**0.250**<br>**0.562**<br>**0.312** |
| **6JDV_B** | **0.447**<br>**0.447** | **0.429**<br>**0.429** | **4RZD_A** | **0.448**<br>**0.360** | **0.394**<br>**0.273** |

| | | | | | |
|---|---|---|---|---|---|
| | 0.681<br>0.681 | 0.653<br>0.653 | | 0.517<br>0.440 | 0.455<br>0.333 |
| 4PQV_A | 0.238<br>0.250<br>0.238<br>0.250 | 0.227<br>0.227<br>0.227<br>0.227 | 4RUM_A | 1.000<br>1.000<br>0.967<br>0.967 | 0.968<br>0.968<br>0.935<br>0.935 |
| 4FRN_A | 0.833<br>0.667<br>0.600<br>0.750 | 0.588<br>0.588<br>0.529<br>0.529 | 3LWO_D | 1.000<br>1.000<br>1.000<br>1.000 | 0.842<br>0.842<br>0.842<br>0.842 |
| 6QN3_A | 1.000<br>1.000<br>0.091<br>0.091 | 0.688<br>0.688<br>0.062<br>0.062 | 6TFE_A | 0.810<br>0.895<br>0.526<br>0.476 | 1.000<br>1.000<br>0.588<br>0.588 |
| 4L81_A | 0.774<br>0.889<br>0.839<br>0.963 | 0.750<br>0.750<br>0.812<br>0.812 | | | |
| **Mean Sensitivity (Recall):** 0.681 ± 0.235, 0.721 ± 0.264<br>0.693 ± 0.229, 0.759 ± 0.239<br>**Mean PPV (Precision):** 0.629 ± 0.256, 0.596 ± 0.278<br>0.634 ± 0.235, 0.607 ± 0.247 | | | | | |
